# Chromatin accessibility analysis uncovers regulatory element landscape in prostate cancer progression

**DOI:** 10.1101/2020.09.08.287268

**Authors:** Joonas Uusi-Mäkelä, Ebrahim Afyounian, Francesco Tabaro, Tomi Häkkinen, Alessandro Lussana, Anastasia Shcherban, Matti Annala, Riikka Nurminen, Kati Kivinummi, Teuvo L.J. Tammela, Alfonso Urbanucci, Leena Latonen, Juha Kesseli, Kirsi J. Granberg, Tapio Visakorpi, Matti Nykter

## Abstract

Aberrant oncogene functions and structural variation alter the chromatin structure in cancer cells. While gene regulation by chromatin states has been studied extensively, chromatin accessibility and its relevance in aberrant gene expression during prostate cancer progression is not well understood. Here, we report a genome-wide chromatin accessibility analysis of clinical tissue samples of benign prostatic hyperplasia (BPH), untreated primary prostate cancer (PC) and castration-resistant prostate cancer (CRPC) and integrative analysis with transcriptome, methylome, and proteome profiles of the same samples to uncover disease-relevant regulatory elements and their association to altered gene expression during prostate cancer progression. While promoter accessibility is consistent during disease initiation and progression, at distal sites chromatin accessibility is variable enabling transcription factors (TFs) binding patterns that are differently activated in different patients and disease stages. We identify consistent progression-related chromatin alterations during the progression to CRPC. By studying the TF binding patterns, we demonstrate the activation and suppression of androgen receptor-driven regulatory programs during PC progression and identify complementary TF regulatory modules characterized by e.g. MYC and glucocorticoid receptor. By correlation analysis we assign at least one putative regulatory region for 62% of genes and 85% of proteins differentially expressed during prostate cancer progression. Taken together, our analysis of the chromatin landscape in PC identifies putative regulatory elements for the majority of cancer-associated genes and characterizes their impact on the cancer phenotype.

## Introduction

Prostate cancer (PC) is a common malignancy with heterogeneous phenotypes in men. In 18% of patients, disease progresses to lethal castration-resistant prostate cancer (CRPC) (Siegel et al., 2018). Recurrent genomic alterations in primary and metastatic PC have been identified and their role in disease progression has been studied extensively (Armenia et al., 2018; Espiritu et al., 2018; Grasso et al., 2012; Gundem et al., 2015; Peng et al., 2015; Quigley et al., 2018; Robinson et al., 2015). In addition to implicating cancer genes, genome sequencing studies have revealed structural variation in non-coding regions, including enhancer elements driving oncogene expression (Takeda et al., 2018; Viswanathan et al., 2018). Epigenetic characterization studies have further extended understanding of the non-coding genome by revealing the role of DNA methylation patterns (Bedford and van Helden, 1987; Börno et al., 2012; Friedlander et al., 2012; Jimenez et al., 2000; Lee et al., 1997; Mahapatra et al., 2012; Varambally et al., 2002; Xu et al., 2012; Zhao et al., 2020), specific transcription factor (TF) binding sites and histone modifications, including the characterization of the active enhancer landscape in PC tissues (Kron et al., 2017; Pomerantz et al., 2015, 2020; Stelloo et al., 2018; Urbanucci et al., 2012, 2017; Yu et al., 2010). Still, how the chromatin landscape evolves during PC progression and drives aberrant transcriptome (Cancer Genome Atlas Research Network, 2015) and proteome (Latonen et al., 2018; Sinha et al., 2019), is unclear.

Genomic aberrations and epigenetic regulation alter chromatin structure in cancer cells (Flavahan et al., 2017; Losada, 2014). Different chromatin accessibility analysis methods have been used to identify the chromatin landscape across cell lines (Thurman et al., 2012), tissues (Roadmap Epigenomics Consortium et al., 2015), and, most recently, tumor tissues (Corces et al., 2018). In PC, the study by Corces et al. uncovered chromatin accessibility changes at single-nucleotide polymorphism that are associated with increased PC susceptibility and illustrated androgen receptor (AR) binding site enrichment in regulatory regions specific to primary PC (Corces et al., 2018). A recent epigenetic study further demonstrated an association between prostate lineage-specific regulatory elements and PC risk loci and somatic mutation density in different stages of PC (Pomerantz et al., 2020). Binding of AR prominently occurs at distal regulatory elements (Massie et al., 2011; Yu et al., 2010), and AR-driven regulatory programs are context-dependent (Sharma et al., 2013; Wang et al., 2009)(Pomerantz et al., 2020)(Sharma et al., 2013; Wang et al., 2009). In PC cells, AR (Urbanucci et al., 2012; Yu et al., 2010), FOXA1 (Adams et al., 2019; Parolia et al., 2019; Sahu et al., 2011), HOXB13 (Chen et al., 2018; Pomerantz et al., 2015), ERG, and CHD1 (Augello et al., 2019) have emerged as epigenetic drivers of disease (Stelloo et al., 2018). More specificly, ERG fusion-positive tumors have a cis-regulatory landscape that is distinct from other tumors (Kron et al., 2017), and aberrant ERG expression has been shown to alter chromatin conformation and regulation in prostate cells (Rickman et al., 2012; Sandoval et al., 2018; Yu et al., 2010).

To gain insight into the role of the chromatin dynamics in determining phenotypes in PC progression, we analyzed chromatin accessibility in a cohort of clinical patient samples of human PC from benign prostatic hyperplasia (BPH), untreated primary prostate cancer (PC), and locally recurrent castration-resistant prostate cancer (CRPC). By integrating DNA, RNA, protein, and DNA methylation data (Annala et al., 2015; Latonen et al., 2018; Ylipää et al., 2015) from the same samples, we provide a comprehensive catalogue of chromatin-related alterations in PC development and progression. Our results highlight high heterogeneity of regulatory elements utilization, complementarity of chromatin accessibility with DNA methylation, and extensive chromatin-driven reprogramming of the AR activity. In this study, we uncover putative regulatory elements for 65% and 85% of progression-related genes and proteins, respectively.

## Results

### ATAC-seq data from human prostate tissues

To study the chromatin landscape’s role in PC development and progression, we first optimized the assay for transposase-accessible chromatin using a sequencing (ATAC-seq) protocol (Buenrostro et al., 2013) for frozen tissue samples. We characterize chromatin accessibility in 11 BPH, 16 PC, and 11 CRPC prostate tissue samples (see **Methods, Supplementary Table 1**). In earlier studies, we have analyzed these same samples using DNA, RNA, and DNA methylation sequencing and SWATH proteomics (**Supplementary Table 1**) (Latonen et al., 2018; Ylipää et al., 2015). Here, these data types were integrated with the ATAC-seq data (**Figure 1A**). ATAC-seq data depth varied from 69 to 204 million reads per sample. Quality control illustrated that there was no significant association between the sequencing depth and key quality parameters such as transcription start site (TSS) enrichment or number of detected peaks (**Supplementary Figure 1A-C**). On the contrary, we observed a good correlation between high quality autosomal alignments (HQAA) and TSS enrichment, indicating a good signal to noise ratio.

**Figure 1:**
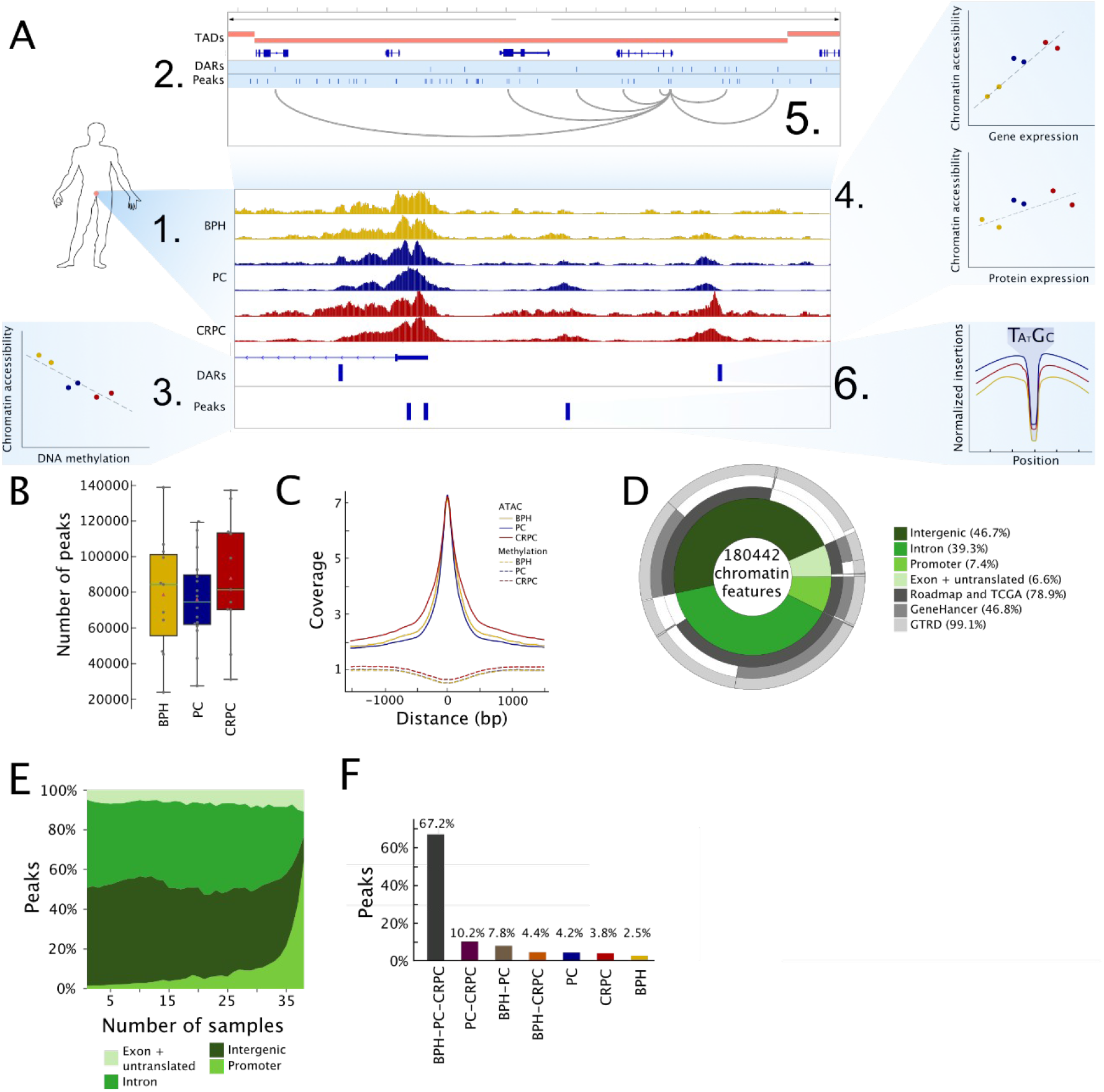
Chromatin accessibility in promoters is robust during prostate cancer progression. **A.** Cartoon illustration of ATAC-seq data analysis. After (1) generating ATAC-seq data from human prostate tissues, we (2) identified peaks and differentially accessible regions (DARs) between BPH, PC and CRPC groups. We (3) compared chromatin accessibility to DNA methylation and (4) gene and protein expression. Next, we associated (5) accessible chromatin regions with correlating target genes within the same topologically associating domains (TADs). Finally, (6) transcription factor binding at accessible chromatin was analyzed using TF footprinting, integration with ChIP-seq data, and using deep learning models to uncover binding context. **B.** Boxplots of the number of raw peaks in each sample (grey dots) in BPH, PC, and CRPC groups are shown. Peak counts in each group are comparable. **C.** Background-corrected coverages from ATAC-seq data at peak locations show a strong signal. Background-corrected DNA methylation data in the same locations is slightly depleted. Distances are relative to peak center. Median signals from BPH, PC, and CRPC samples are shown. **D.** Chromatin features are ordered in the donut plot based on their annotation to genomic location categories: intergenic, intron, promoter, and exon and untranslated regions (5’-UTR, 3’-UTR, transcription termination sites, non-coding RNA). Majority of the features are located in intergenic and intronic regions. For each category, the proportion of previously identified areas of accessible chromatin (Roadmap and TCGA), known enhancer regions (GeneHancer), and detected TF binding sites from ChIP-seq data (GTRD) are shown. Fraction of chromatin features belonging to each region is shown in the donut plot with percentages given in labels. **E.** The proportion of peaks located in genomic location categories is shown for peaks present in the different number of samples. Most consistently observed ATAC peaks are located at promoters, and peaks in the distal regions are more heterogeneous across the samples. **F.** Percentage of peaks in different sample group combinations. Although the majority of peaks are present in samples from all three sample groups, a subset of peaks are sample group-specific.

### Chromatin accessibility at distal sites is heterogeneous in prostate cancer

To identify accessible and progression-related chromatin features, we used two complementary approaches. In the first approach, accessible chromatin regions in each sample were identified by peak calling using MACS2 peak calling algorithm (Zhang et al., 2008). We identified 23,840 to 138,942 raw peaks per sample (**Supplementary Table 1**). The number of detected peaks was not characteristic to a specific sample group, but samples with high and low peak count were observed throughout BPH, PC, and CRPC groups (**Figure 1B**). To obtain a robust set of reproducible peaks across samples, we used a previously proposed approach to unify raw peak calls (see **Methods**)(Corces et al., 2018). This approach resulted in the compilation of 178,206 peaks across the sample set (**Supplementary Table 1**). This is consistent with previous estimates for the number of cancer type-specific peaks in chromatin accessibility data (Corces et al., 2018). In the second approach, we performed genome-wide analysis to identify differentially accessible regions (DARs) by comparing samples in BPH to PC and PC to CRPC groups (see **Methods**). As a result, we identified 1,727 and 3,498 differentially accessible regions (DARs) for BPH to PC and PC to CRPC, respectively, with false discovery rate (FDR) below 10% (**Supplementary Table 2**). For peaks and DARs, a clear chromatin accessibility signal is detected (**Figure 1C**, **Supplementary Figure 2A**) and DNA methylation is depleted (**Figure 1C**, **Supplementary Figure 2A**) consistent with previous studies reporting decreased DNA methylation at accessible chromatin loci (Corces et al., 2018; Urbanucci et al., 2017).

Of the 180,442 identified chromatin features, 72% overlapped with regulatory regions found in normal tissues (Corces et al., 2018; Roadmap Epigenomics Consortium et al., 2015) or TCGA data (Corces et al., 2018; Roadmap Epigenomics Consortium et al., 2015) (**Figure 1D**). The overlap was consistent for both peaks and DARs (**Supplementary Figure 2B**). TCGA data included 20 primary PC samples and within these samples 65.8% of their peaks overlap with our peak set. Taken together, our data showed consistency with earlier chromatin accessibility studies and we were able to expand the known regulatory landscape by discovering 38,157 new prostate cancer related chromatin features.

Of all identified chromatin features 7.4% were in promoters, 6.6% were in exons and untranslated regions, 39.3% were in introns (51.8% overlapping previously marked enhancers), and 46.7% were intergenic (32.5% overlapping previously marked enhancers) (Fishilevich et al., 2017) (**Figure 1D**). Peaks and DARs were distributed similarly, except for the promoter region in which 7.4% of the peaks but only 1.9% to 2.4% of the DARs were located (**Supplementary Figure 2B**). Furthermore, the peaks located at promoters had higher signal intensity than peaks in other genomic annotation groups (**Supplementary Figure 2C-D**). In addition, 60% of the peaks common to all the samples are located on promoters (**Figure 1E**, **Supplementary Figure 2E**). When assigning the peaks to a sample group or groups based on if they are present in a specific sample (**Figure 1F**), we observed that most peaks are not group-specific. For the peaks assigned to each sample group, the annotation distribution is similar (**Supplementary Figure 2F**). Importantly, we did not observe any peaks that would be group-specific and present in all the samples of that group (**Supplementary Figure 2G**). These data show that while promoters are robustly open across samples, accessibility at other genomic regions is highly variable between samples and sample groups. This indicates that, while accessibility remains robust during PC progression, most of the chromatin alterations occur at intronic and intergenic regions.

### Progression-related chromatin alterations are consistent

Having characterized chromatin features, we looked into its alterations over disease progression. Comparing DARs in BPH to PC and PC to CRPC we found little overlap (**Figure 2A**) suggesting that differential accessibility-related chromatin changes are specific to PC initiation and to progression of CRPC (**Figure 2B**, **Supplementary Figure 3A**). DARs in PC to CRPC comparison show a clear increase in untranslated and exon regions (**Supplementary Figure 2B, Supplementary Figure 3B**). These loci are not usually reported to harbor gene regulatory elements, but this combined with the finding that CRPC samples show more opening DARs than the other group (**Figure 2A**, **Supplementary Figure 2B**) may reflect overall chromatin relaxation (Urbanucci et al., 2017) or events related to chromatin reorganization.

**Figure 2:**
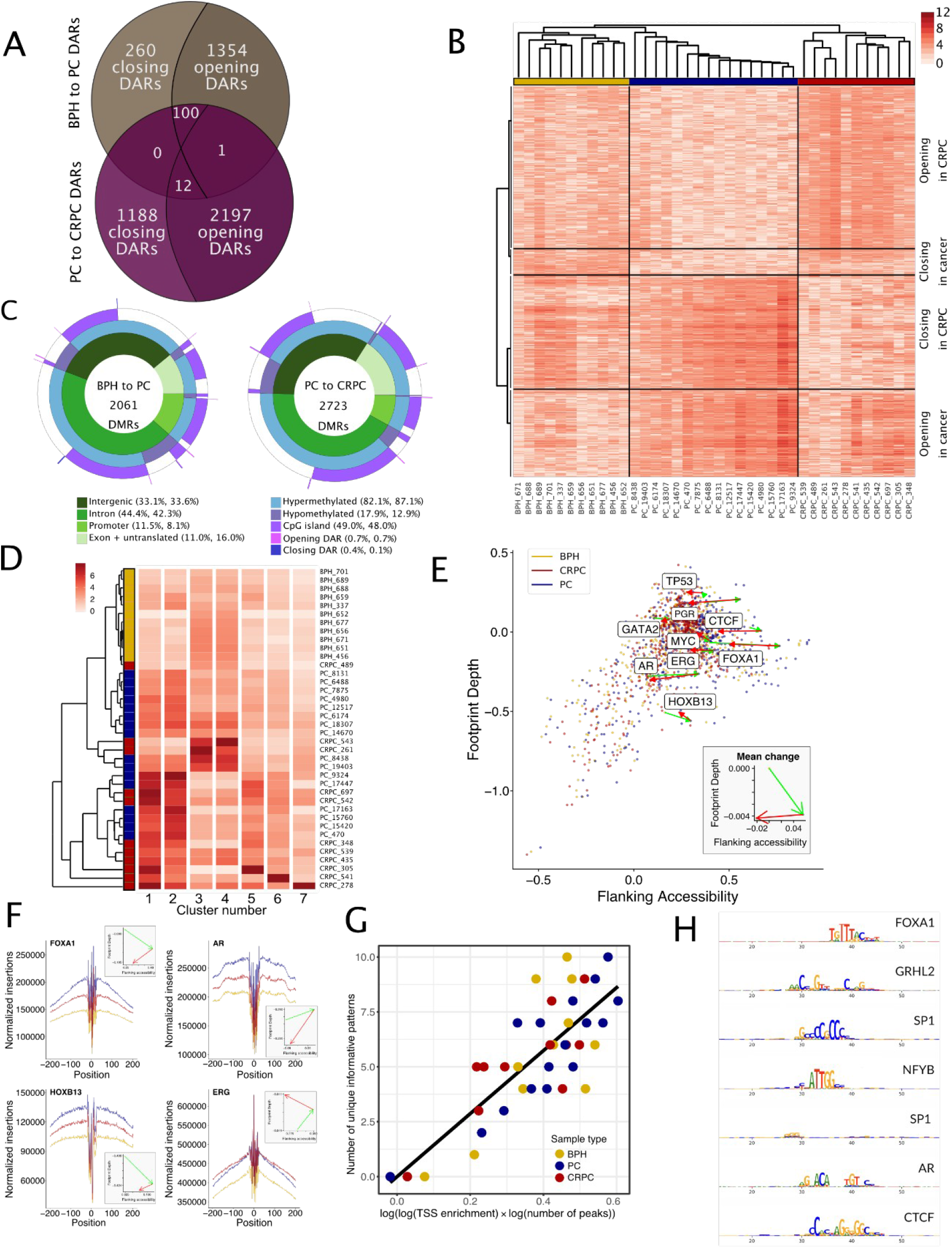
Differential accessibility is concentrated on regulatory regions. **A.** Venn diagram showing the numbers BPH to PC and PC to CRPC DARs and their overlap. Only a small portion of DARs are shared between comparisons. **B.** Clustering of samples using ATAC-seq signal of DARs separates them into BPH, PC, and CRPC groups, and identifies progression-related chromatin accessibility patterns. Scale bar shows log2 of normalized ATAC-seq signal. Pearson correlation was used as the distance metric, and linkage was calculated using the Weighted Pair Group Method with Arithmetic Mean (WPGMA) algorithm. **C.** Donut plots show genomic location categories for DMRs from BPH to PC and PC to CRPC comparison groups. Rings show whether DMR is hypermethylated or hypomethylated and whether it is located in a known CpG-island (within +/2kb). The outermost wedges show overlap with opening and closing DARs. In labels, percentages are given for BPH to PC and PC to CRPC DMRs, respectively. Differential accessibility and DNA methylation during progression occur at distinct loci. **D.** Unsupervised clustering of cancer-specific peaks shows clear clusters but fails to separate PC samples from CRPC samples. **E.** TF footprinting based on Tn5 transposase insertion sites was done for all expressed TFs with HOCOMOCO motif to quantify flanking accessibility and footprint depth. Averages from BPH, PC, and CRPC samples are shown and transitions in footprinting space (BPH to PC in green, PC to CRPC in red) are illustrated for PC-related TFs and those with the largest change between groups. Mean change is shown in the inset. **F.** Detailed TF footprints for key TFs AR, FOXA1, HOXB13, and ERG to illustrate the change in chromatin accessibility during progression. Quantification of footprint depth and flanking accessibility are shown in the insets. **G.** Motif discovery with BPNET correlates with the signal to noise (TSS enrichment) and the number of peaks used in training. **H.** Example of discovered motifs with BPNET on high quality sample PC_9324.

Using methylated DNA immunoprecipitation sequencing (MeDIP-seq) data on the same clinical samples, we also called progression-related differentially methylated regions (DMRs) (see **Methods**). Comparing BPH to PC and PC to CRPC, we found 2,061 and 2,723 DMRs (**Supplementary Table 2,** **Figure 2C**). Comparing DARs and DMRs, we detected only 13 (0.6%) and 23 (0.8%) overlapping features in each comparison(**Figure 2C**, **Supplementary Figure 2B, Supplementary Figure 3C, Supplementary Table 2**). Little overlap between DARs and DMRs suggests that regulation of chromatin accessibility and DNA methylation might work as distinct epigenetic regulatory mechanisms in PC, affecting different transcriptional outputs.

### Heterogeneity in chromatin accessibility is associated with disease-relevant regulators

As most of the observed chromatin alterations occur at intronic and intergenic regions, to understand how the heterogeneity of chromatin relates to disease progression, we first focused on the cancer-specific peaks (**Figure 1F**) with highest variance in signal across the samples. This includes mostly peaks distal from TSS, whilst promoter peaks are depleted in this set (**Supplementary Figure 3D)**. Unsupervised analysis of these peaks (see **Methods**) separated the samples in three clusters containing 273 to 1655 peaks, but failed to separate PC and CRPC samples in a data-driven manner (**Figure 2D**, **Supplementary Figure 3E)**. The three clusters did not correlate with tumor class/state, Gleason score, or ERG fusion status. However, the peaks separated in seven clusters based on consensus clustering (**Supplementary Figure 3F**). Enrichment analysis using TF binding site predictions in each of the seven peak clusters was used to evaluate whether these contained regulatory regions for specific TFs (**Supplementary Figure 3E, Supplementary Table 1**). Interestingly, each peak cluster is associated with DNA binding of different PC-related TFs. ERG-enriched and AR- and FOXA1-enriched clusters showed a similar activity pattern across samples. Likewise epithelial to mesenchymal transition (EMT) associated Wnt/β-Catenin signaling and TEAD1 and SNAI1 clusters behave similarly (Odero-Marah et al., 2018; Zhou et al., 2016). AR pioneering factors GATA2 and HOXB13 (Hankey et al., 2020; Pomerantz et al., 2015) were enriched into the same cluster, which showed the highest accessibility in the CRPC-rich sample group. Other clusters represent sample specific signals, for example, immune response related TFs were highly accessible only in one CRPC sample, possibly due to the patient’s immune response.

To further study the effect of chromatin accessibility variation on TF activity, we performed TF footprint analysis in each sample for expressed TFs with available binding motif (see **Methods**, **Figure 2E**, **Supplementary Table 3**). Quantification of TF footprint by “flanking accessibility” (FA) and “footprint depth” (FD) allows the study of TF activities in a genome-wide manner (Baek et al., 2017). For the majority of TFs, FA and FD correlate. Notably, we do not detect any TF e.g. with low FA and high FD. Several disease relevant TFs, including AR and FOXA1, are among the ones with largest change in FA and FD during progression (**Figure 2E**). AR, and related co-factors FOXA1, and HOXB13 have similar Tn5 insertion patterns with the highest accessibility in PC (**Figure 2F**) while ERG accessibility is similar in all sample groups. During progression to CRPC, CTCF displays a large change in FA which might reflect relaxation or other alterations of chromatin structure.

Taken together, these results highlight a highly heterogeneous chromatin landscape across samples, and demonstrate that the observed regulatory patterns are associated with known disease-relevant processes and regulators. Furthermore, changes in disease relevant TF activities are consistent over progression.

### Similar TF binding syntaxes are conserved across tumor samples

To understand if the heterogeneity in chromatin accessibility leads to variability in TF binding syntax, we utilized the recently developed BPNET model (Avsec et al.)(see **Methods**). BPNET builds predictive models of chromatin accessibility, and recursively decomposes the output to assign base-pair contribution scores to every input sequence that can be combined to obtain binding syntax motifs. We tested the model with cell line data and were able to discover highly detailed binding patterns, e.g. different forms of known AR binding configuration, demonstrating the feasibility of the approach with ATAC-seq data (**Supplementary Figure 4**). With application to data from patient samples we observed that model performance is dependent on both the signal-to-noise ratio and the number of available training peaks (**Figure 2G**). When we trained the model on individual samples using the whole reproducible peak set, we were able to recover motifs that match with known TFs, including AR, FOXA1, CTCF, GRHL2 and SP family (**Figure 2H**, **Supplementary Table 3**). Despite high heterogeneity in peaks across samples, detected binding syntaxes are consistent. When performing model training using only peaks with a known DNA binding site (see **Methods**) for key TFs AR, FOXA1, or HOXB13, we observed a consensus binding motif for all tested TFs across high quality samples (**Supplementary Table 3, Supplementary Figure 5-8**). This observation further supports the idea that TFs binding properties do not change despite heterogeneity in chromatin accessibility. In addition, we were able to identify disease state-related factors co-occurring with selected driver TFs (**Supplementary Table 3, Supplementary Figure 5-8**).

### Distal regulatory elements accessibility correlate with expression of disease relevant genes

To gain insight into the functional role of accessible chromatin, we integrated ATAC-seq data with RNA and protein expression data from the same samples (see **Methods**). While promoter accessibility was consistent across samples, correlation between gene expression and transcription start sites (TSS) accessibility is very moderate (Spearman correlation ρ = 0.11 and ρ = 0.04 for RNA and protein data, respectively; **Figure 3A**, **Supplementary Table 4**). Analysis of gene groups with different expression levels (high, moderate, low, and housekeeping genes) suggests that this is due to promoters of expressed genes being mostly open in basal state (**Figure 3B**). However, differential chromatin accessibility at TSS and differential expression between groups are still co-occurring. For differentially expressed (D.E.) genes in the BPH to PC comparison, we observed an enrichment of genes with association between accessibility and expression (Fisher’s exact test p < 10^-16^, **Figure 3C**). These included several PC-related oncogenes such as *AR*, *MYC*, and *BCL11A* (**Figure 3D**). In the PC to CRPC comparison, there was an enrichment of genes in which TSS closing was associated with decreased expression (Fisher’s exact test p = 9.19 * 10^-16^, **Figure 3C**). Overall, from the promoter-proximal regions (−1kbp/+100bp), we detected 418 peaks, one BPH to PC DAR, and 9 PC to CRPC DARs with strong correlation (|correlation coefficient| > 0.5) to expression for the adjacent gene (**Supplementary Table 4**). For the PC to CRPC comparison, eight out of nine DARs showed increased accessibility. The remainder DAR shows reduced accessibility in CRPC and is located in the promoter of the *MIR30A* gene, which codes for a tumor suppressor miRNA (Jiang et al., 2018) downregulated in CRPC (log2 fold change −1.2389, Spearman ρ = 0.7 p = 6.18*10^-5^). Thus, while global correlation is moderate, the expression of several disease-relevant genes is strongly correlated with promoter accessibility, suggesting reconfiguration of the promoter state during disease progression.

**Figure 3:**
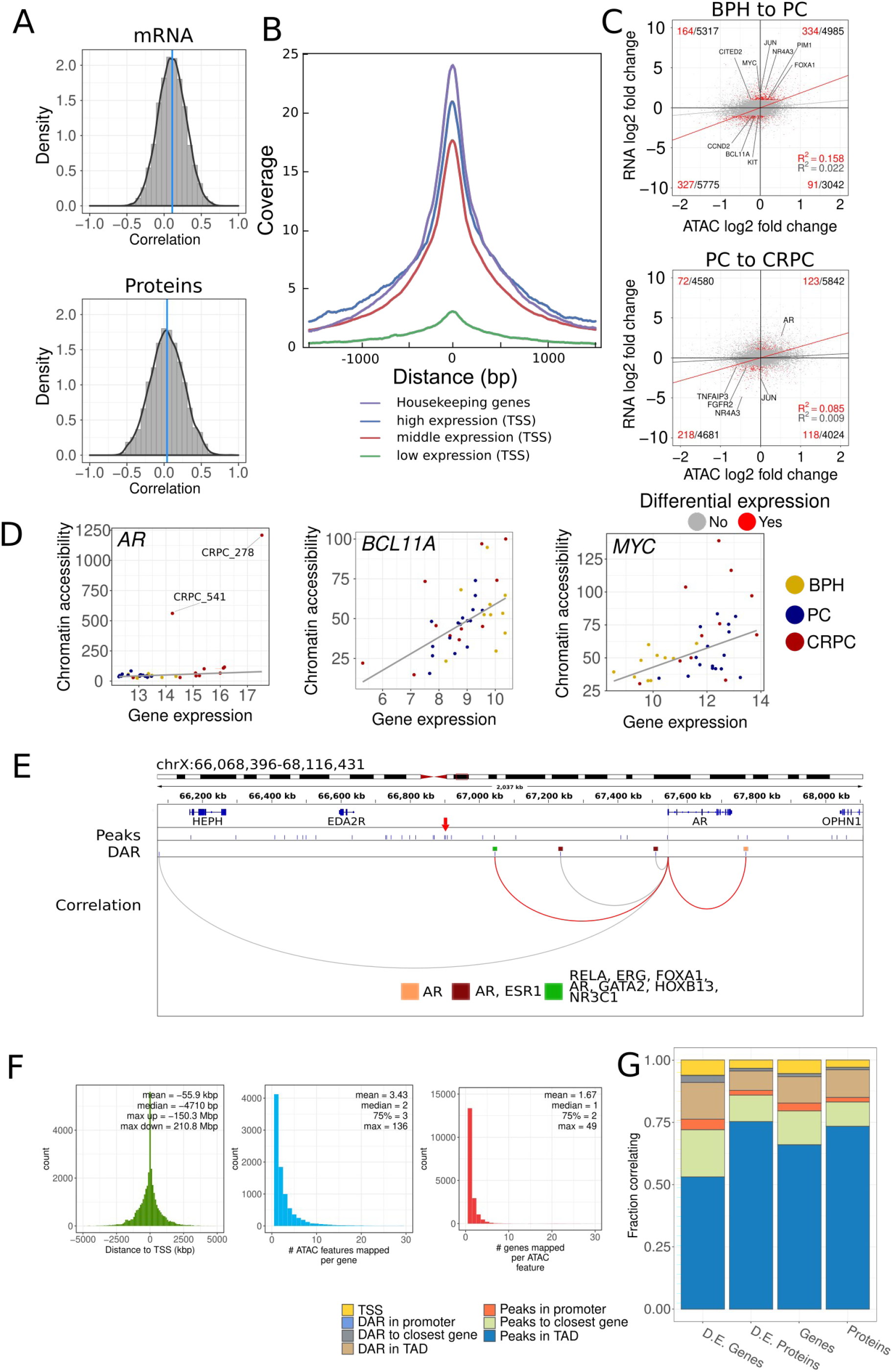
Distal features detected by ATAC-seq are correlated with gene expression in prostate cancer samples. **A.** Correlation between TSS chromatin accessibility and gene expression is moderate at the genome-wide scale. Density plot of Spearman correlation coefficient between gene (top, median=0.11) or protein (bottom, median=0.04) expressions and normalized ATAC-seq signal at the TSS. **B.** ATAC-seq background-corrected coverage on TSS. TSSs are grouped based on expression of the gene (high, middle, and low (see **Methods**)) or annotation to housekeeping genes. Chromatin of expressed genes is accessible at TSS. Low expression genes show minimal chromatin accessibility. **C.** Differential expression and chromatin accessibility have positive association. Scatter plots visualize the association between differential RNA expression and TSS accessibility in BPH to PC (top panel) and PC to CRPC (bottom panel) comparisons. Differentially expressed genes are shown in red. Gray and red lines show regression lines fitted to their corresponding data points to demonstrate the association between data types. Selected oncogenes are labeled. Numbers in the corners of each quadrant of the scatter plot report counts of differentially expressed and total genes. Differentially expressed genes are enriched for opening chromatin and increased expression, and closing and decreased expression in BPH to PC comparison. In PC to CRPC comparison, enrichment is seen only in closing and decreased expression quadrant. **D.** Correlation between chromatin accessibility and gene expression for the selected oncogenes demonstrate increasing (*AR, MYC*) and decreasing (*BCL11A*) accessibility during progression. For *AR*, outlier chromatin accessibility is observed for samples with high-level amplification identified from DNA-seq data (CRPC_278, CRPC_541). **E.** Correlation analysis between chromatin accessibility and gene expression identifies putative regulatory elements. In total 48 peaks and 5 DARs are detected in a 2 Mbp TAD region around the *AR* locus. Known associations from GeneHancer database are shown in red. Binding sites for selected TFs from GTRD database within associated DARs are shown. Red arrow indicates a peak detected at recently reported *AR* enhancer locus. **F.** Characterization of correlations shows that associations between regulatory elements and genes are specific. Left panel shows the distance of correlating chromatin features from TTS. Middle panel indicates the number of chromatin features mapped to each gene. Finally, the last panel gives the number of genes mapped to each chromatin feature. Summary statistics are given in the insets. Mean, median, and maximum upstream (max up) and downstream (max down) distances are reported for the distance distribution. For the middle and right panels, mean, median, upper quartile and maximum number of associations are reported. **G.** Summary of all correlation analyses. Fraction of genes and proteins correlating with ATAC-seq features across all analyses is reported. Data for all and differentially expressed gene subsets are shown.

Next, we focused on understanding how distal accessible chromatin sites, that vary the most across the sample set, associate with gene expression changes. Co-regulated genes are found within topologically associating domains (TAD, (Pombo and Dillon, 2015)). Therefore, to limit the target gene associations to a biologically meaningful context, we used previously published annotations of TADs (Pombo and Dillon, 2015) from PC cells (see **Methods**). Within these TADs boundaries, we identified all peak-gene and DAR-gene pairs with a strong correlation (see **Methods**, **Supplementary Figure 9**). All together 9.6% (17,066) of all peaks and 25.4% (1300) of DARs were assigned to putative target genes based on correlation (**Supplementary Table 4**), including 8977 unique genes from 1871 TADs. We found that 29.6% of PC to CRPC DARs correlate with gene expression while only 16.4% of BPH to PC DARs correlate. Ingenuity Pathway Analysis (IPA) performed separately for genes associated with either DARs or peaks showed several PC-related and cancer-related pathways enriched (**Supplementary Table 4**), demonstrating that chromatin-related changes reflect disease-relevant target gene alterations.

When looking into associations with specific PC genes, we found 5 PC to CRPC DARs and 48 peaks with strong correlation to *AR* expression (**Figure 3E**, **Supplementary Table 4**). DARs correlated with *AR* expression are located within 2 Mbp region around the *AR* locus, indicating regulatory potential throughout the TAD area. These DARs harbor binding sites for key TFs including *AR*, *FOXA1*, *HOXB13*, and *ERG* (**Figure 3E**). Peaks correlating with *AR* expression are mainly upstream of TSSs (41/48) (**Figure 3E**). Identified peaks include an enhancer known to be amplified in advanced PC ((Takeda et al., 2018; Viswanathan et al., 2018)). The expression of 42 known oncogenes (e.g. *EGFR, ERBB2, JUN, FGFR1* and *FGFR2*), 27 tumor suppressor genes (e.g. *NOTCH1, BRCA1, BRCA2, IL2*) and 22 genes related to chromatin regulation (e.g. *HDAC1, HDAC2, HDAC5, HDAC6, HDAC9, HDAC10,* and *SMARCD1*) correlated with the chromatin accessibility of at least one peak (**Supplementary Table 4**). In addition, the expression of 4 oncogenes (*JUN, PIM1, CARD11,* and *TFG*), 5 tumor suppressor genes (*PTEN, NOTCH1, CDK6, FH,* and *WT1*) and 2 factors involved in chromatin regulation (*HDAC7* and *CHRAC1*) were strongly associated with DARs irrespective of the comparison group (**Supplementary Table 4**).

Analyses of distal and promoter areas identified 418 associations from TSS signal to gene expression as well as 27,353 peak–gene and 3,513 DAR-gene pairs. Expression-associated areas of chromatin accessibility are mostly located close to TSSs (median distance 4.7 kbp upstream of TSS) (**Figure 3F**). Also, 45.8% (4,124) of genes with expression correlating with chromatin accessibility are linked to exactly one regulatory element, while 97 genes (1.07%) can be associated to 30 or more regulatory elements (mean= 3.4, **Figure 3F**, **middle**). Likewise, 72.4% (13,359) of peaks or DARs correlating with gene expression are associated with a single gene and 35 are linked to 30 or more unique genes, indicating that those might be regulatory hubs (mean=1.7, **Figure 3F**, **right**). Taken together, we could associate at least one peak or DAR to 45.5% of genes and 30.8% of proteins (**Figure 3G**, **Supplementary Table 4**). When focusing on the genes with differential expression patterns, 62.4% and 84.7% of genes and proteins were associated, respectively. As reported earlier, correlations at the transcript and protein levels are not consistent (Latonen et al., 2018; Sinha et al., 2019), but both data levels support the conclusion that the majority of differential expression in progression-related genes can be correlated with chromatin accessibility.

### Chromatin accessibility alterations during disease progression are associated with different transcription factors regulatory modules

To gain understanding on how the chromatin accessible sites direct transcriptional programs during PC progression, we generated TF–gene expression regulatory network. TFs were connected to their target genes through known binding sites in accessible chromatin regions (see **Methods**). We focused this analysis specifically on DARs that correlate with gene expression (**Figure 4A**). From the TF-gene network that we generated, we identified regulatory modules, defined as a set of TFs that share a set of target genes (see **Methods**). Two clear modules with 1082 and 799 target genes emerged from the analysis. The module with the largest number of target genes represents the well-characterized AR regulatory program, including AR, FOXA1, and ERG (**Figure 4B**, **Supplementary Figure 10A**). The second module contains a number of TFs with known function in driving aggressive prostate cancer e.g. glucocorticoid receptor (NR3C1) as well as TF coding genes MYC, HOXB13, GATA2, NKX3-1, and PGR (Chen et al., 2018; Grindstad et al., 2018; Isikbay et al., 2014; Koh et al., 2010; Rodriguez-Bravo et al., 2017). Surprisingly, genes targeted by this second module are a subset of AR module target genes (**Figure 4C**). We validated this by repeating the analysis using peaks instead of DARs (**Figure 4D**, **Supplementary Figure 10B-D**). IPA analysis of target genes confirmed AR as an upstream regulator for both modules (**Supplementary Table 4**), but in the second module, AR activity is predicted to be inhibited. This suggests that this second TF module could compensate for reduced AR activity e.g. due to androgen deprivation treatment. This was clearly shown for glucocorticoid receptor which is upregulated in CRPC especially resistant to enzalutamide treatment (Arora et al., 2013).

**Figure 4:**
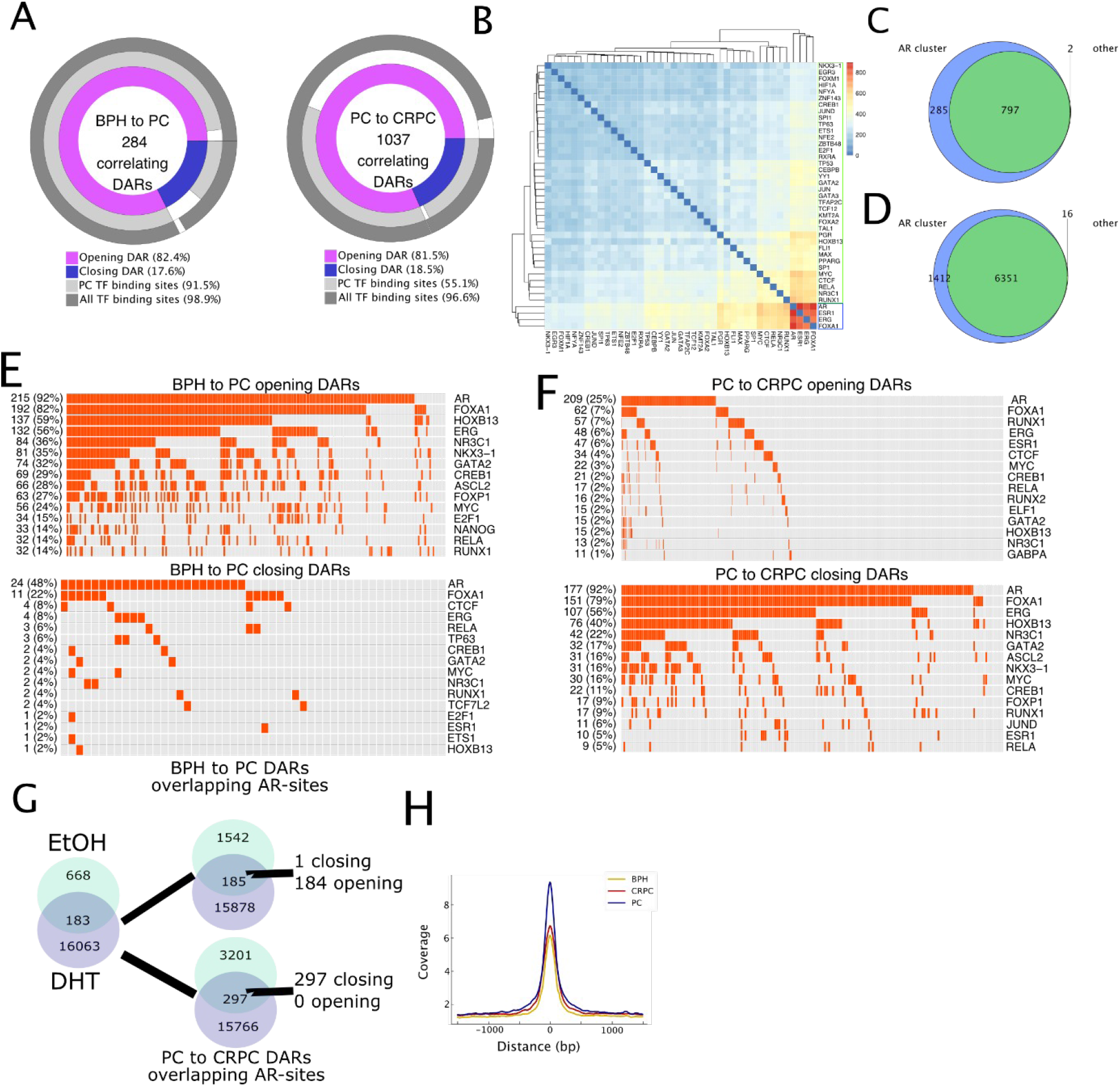
Disease progression alters prostate cancer-specific transcription factor binding site accessibility and regulatory programs. **A.** Donut plots showing numbers of gene expression correlating DARs in BPH to PC (left) and PC to CRPC (right) comparisons. Shown are also percentages of opening and closing sites and whether they harbour TF binding sites as characterized in the GTRD database. **B.** Hierarchical clustering of TF gene expression network uncovers two groups of TFs: a core cluster composed of AR, ERG, FOXA1 and ESR1, and a second cluster sharing a high number of target genes with the AR core cluster. Complete linkage and euclidean distance were used in clustering. Scale bar encodes the number of shared genes. **C**. Venn diagram shows that the two TF clusters indicated in B share a substantial amount of target genes. **D.** Repeating the intersection analysis with genes linked to peaks, a similar pattern as in C is observed. **E.** Oncoprints illustrate 15 TFs with the highest number of binding sites (taken from GTRD prostate cancer subset) overlapping with gene expression correlating DARs. Panels represent sites from BPH to PC opening (top) and closing (bottom) DARs. AR binding sites are present in almost all (92%) opening sites in this comparison. **F.** Similar oncoprints as in E but for PC to CRPC opening (top) and closing (bottom) DARs. In this comparison, most of the closing sites (92%) include AR binding sites **G.** Androgen-induced AR binding sites taken from Massie et al. 2011 within DARs are present in opening regions in BPH to PC comparison and in closing regions in PC to CRPC comparison. **H.** Background-corrected ATAC-seq coverage of AR binding sites from androgen-treated (DHT) cells (Massie et al. 2011) is stronger in PC samples than other sample groups.

To elucidate the interplay of TFs in more detail, we performed a comparative analysis of TF binding sites, identified from prostate cancer cells, in opening and closing DARs (**Figure 4A**). In DARs from BPH to PC comparison AR, FOXA1 and HOXB13 binding sites are the most abundant and are co-occurring within 53.1% and 2.1% of opening and closing AR sites, respectively (**Figure 4E**, **Supplementary Figure 10E**). In PC to CRPC DARs, we observed the opposite pattern with 1.6% and 36.1% of opening and closing AR sites, respectively (**Figure 4F**, **Supplementary Figure 10E**). Again, we observed consistent correlations when repeating the analysis using peaks instead of DARs (**Supplementary Figure 5F).** These results suggest that chromatin opening in PC remains mostly accessible also in CRPC and harbour AR binding sites. Moreover, in CRPC new chromatin opening events enable additional TFs to bind the regulatory regions (**Supplementary Figure 10G-H**). Concomitantly, in CRPC several AR binding sites are closing, consistent with reduced AR activity in CRPC samples (p=0.02, **Supplementary Figure 10I**).

To test whether the chromatin in CRPC is selectively closed in AR binding sites related to canonical AR regulation, we used publicly available cell line data (Massie et al. 2011). To study the interplay between AR chromatin binding, androgen stimulation and chromatin accessibility we evaluated the overlap between androgen-induced AR binding sites in cell lines and DARs (**Figure 4G**, **Supplementary Figure 11A**). The majority of DARs are open in BPH to PC and closed in PC to CRPC comparison, which confirms our hypothesis that the canonical AR regulation is suppressed during progression to CRPC. In agreement with this observation, more PC-specific ATAC-seq peaks overlap these AR binding sites than CRPC- or BPH-specific peaks (**Supplementary Figure 11B**). We also note that the AR binding site locations from the cell line have most accessible chromatin in PC samples (**Figure 4H**, **Supplementary Figure 11C-F**).

## Discussion

In this study we integrated for the first time data on chromatin accessibility, DNA methylation, transcriptome, and proteome in clinical BPH, PC, and CRPC tissue samples. We used ATAC-seq to define a catalogue of accessible genomic regions and to characterize changes in chromatin accessibility during PC progression. The identified open chromatin regions are consistent with previous chromatin accessibility studies (Roadmap Epigenomics Consortium et al., 2015); (Corces et al., 2018). Furthermore, the number of detected peaks is consistent with earlier predictions of cancer type-specific peaks (Corces et al., 2018). Our analysis extended the known chromatin landscape by 38,157 reproducible previously uncovered accessible chromatin sites specific for PC. The majority of these sites have previously reported TF binding activity.

The chromatin accessibility of PC shows inter-sample heterogeneity. While we observed consistent accessibility at promoter regions during disease progression, accessibility does not correlate well with gene expression at the genome-wide level. As gene expression is regulated by the repressive or activating functions of the TFs binding to the promoters and distal regulatory elements, it is clear that promoter accessibility signal alone cannot be highly predictive of expression, as reported also by several earlier ATAC-seq studies across different systems (Rajbhandari et al., 2018; Scharer et al., 2018; Toenhake et al., 2018; Wu et al., 2018). This also highlights the important role of enhancers and their regulation in driving tumor development and progression. We did observe strong correlation with promoter or putative enhancer accessibility to gene expression for a subset of PC-related genes. At least one putative accessible regulatory element was found for 62.4% of protein coding genes and 84.7% of proteins with a differential expression. The majority of these regulatory elements are from the peaks and DARs that correlate with genes within the same TAD, providing a rich resource of candidate genes and regulatory elements for future investigation.

Still, a large fraction of putative regulatory regions could not be associated with genes. This might be explained by our utilization of stringent criteria for detecting target genes because of the limited cohort size. In addition, we used predefined TAD structure in the analysis and thus, our analysis could not detect associations resulting from altered TAD boundaries (Taberlay et al., 2016). Furthermore, many of the identified regions might contribute to functions other than direct regulation of gene expression. For example, it is known that higher order chromatin structure alterations may occur in PC tumorigenesis (Gerhauser et al., 2018), such as chromatin compartment formation and looping (Gerhauser et al., 2018; Rowley et al., 2018; Weischenfeldt et al., 2017). We did observe a large number of CTCF binding sites in peaks and DARs that may partially reflect these phenomena. Moreover, the majority of DARs in BPH to PC and PC to CRPC were opening (84.3% and 63.2%, respectively), supporting the idea that chromatin in PC initiation and progression undergoes a process of continued relaxation (Urbanucci et al., 2017)(Braadland and Urbanucci, 2019).

Chromatin accessibility and DNA methylation had the expected inverse relationship at the genome-wide level. The increase in the number of DMRs in the PC to CRPC comparison was not as significant as the two-fold increase in the number of DARs. This indicates that methylation-independent changes in chromatin accessibility are more prevalent during the progression to CRPC. Furthermore, DMRs and DARs overlapped in only a few regions, suggesting that these two epigenetic mechanisms are driving different transcriptional regulatory programs. Earlier work has shown the interplay between chromatin modifications and DNA methylation through interaction of EZH2 with DNA methyltransferases (DNMTs) (Viré et al., 2006). Further studies are needed to better understand how differential regulation of DNA methylation and chromatin accessibility are targeted.

Integration of TF binding data and predictions with accessible chromatin areas allowed us to analyze the regulatory programs that are associated with the identified peaks and DARs. Analysis of TF binding patterns demonstrated that despite high variability in chromatin accessibility, the observed motifs and TF enrichments are consistent during PC evolution. This suggests that there are a number of different chromatin configurations that can lead to similar phenotypes. For instance, AR was identified among the top candidate regulators but at the same time, the AR gene was one of the most targeted genes by chromatin remodelling during PC progression. A number of sites with accessibility were present in the genomic neighborhood of *AR,* including a previously reported AR-enhancer site, which was shown to be activated by structural rearrangement (Takeda et al., 2018). The analysis revealed that the interplay between AR, FOXA1, and HOXB13 TFs (Pomerantz et al., 2015) was the most prominent PC initiation-associated transcriptional regulatory module. FOXA1 is known to pioneer TFs binding to chromatin, including AR (Lupien et al., 2008) (Jozwik and Carroll, 2012). HOXB13 is a prostate lineage-specific TF and germline alterations have been shown to increase PC risk (Ewing et al., 2012). Previous studies with PC cell-lines identified alternative AR programs in CRPC (Sharma et al., 2013; Wang et al., 2009). Here we were able to show that this AR, FOXA1, HOXB13 program is initially activated in PC then depleted during progression to CRPC, when it is substituted by the activation of alternative regulatory modules composed of several TFs previously reported to be important in progression to CRPC. These TFs include glucocorticoid receptor, known to have a role in developing resistance to antiandrogens (Arora et al., 2013), and progesterone receptor that has been associated with disease progression (Grindstad et al., 2015, 2018). Overall, these analyses demonstrate that epigenetic chromatin reprogramming during CRPC progression enables binding sites for disease driving TFs, in addition to AR.

In summary, we demonstrated how transcriptional regulatory programs are altered in PC progression by characterizing the chromatin accessibility landscape and its alterations in human PC tissue. We reveal regulatory elements that are activated in PC and identify putative regulators for known oncogenic and tumor suppressive genes.

## Methods

### Sample collection

Fresh frozen tissue specimens were acquired from Tampere University Hospital (Tampere, Finland). 11 BPH, 16 untreated PC, and 11 CRPC samples were used for ATAC-seq library generation. BPH samples included were collected either by transurethral resection of the prostate (TURP; n=4) or radical prostatectomy (RP; n=7) (**Supplementary Table 1**). PC samples were obtained by radical prostatectomy. Locally recurrent CRPC samples were obtained by transurethral resection of the prostate. Samples were snap-frozen and stored in liquid nitrogen. Histological evaluation and Gleason grading was performed by a pathologist based on hematoxylin/eosin-stained slides. All samples contained a minimum of 70% cancerous or hyperplastic cells. The use of clinical material was approved by the ethical committee of the Tampere University Hospital. Written informed consent was obtained from the donors.

### Tissue sample processing

Samples were cut from the frozen blocks as 2×50 µm sections. Nuclei were isolated from these sections. All the steps were performed on ice. First 6 ml of ice cold lysis buffer (10 mM Tris·Cl, pH 7.4, 10 mM NaCl, 3 mMMgCl2, 0.1% (v/v) Igepal CA-630, 1× protease inhibitors (Roche, cOmplete)) was added to pre-cooled petri dish and sections were moved from tube to petri dish with 1 ml of lysis buffer. Sections were cut into smaller pieces with a scalpel. Buffer and sections pieces were moved to a 15 ml Falcon tube. Each sample was pulled through a 16 G needle 15 times. Larger pieces were let to sink to the bottom. Supernatant was moved into a new tube and centrifuged at 700 g for 10 min at 4 °C. Supernatant was removed and the pellet was dissolved in a PBS buffer. Nuclei were counted and 50,000 nuclei were transferred to a new tube. Nuclei were pelleted by centrifugation at 700 g for 10 min at 4 °C. Supernatant was removed.

### Processing of cell lines

VCaP cells were cultured in culbecco’s modified eagle’s medium with 10% fetal bovine serum and 1% L-glutamine. Cells were harvested using trypsin and counted. We took 50,000 cells and centrifuged them at 500 x g for 5 min, 4°C. Cells were washed once with 50 µl cold 1xPBS buffer and centrifuged again with the same settings. Supernatant was removed and cells resuspended to 50 µl of cold lysis buffer followed by centrifugation with the same settings. Supernatant was removed.

### ATAC-seq library generation and sequencing

ATAC-seq libraries were generated as presented earlier (Buenrostro et al., 2013). Briefly, transposition mix (25 μl 2× TD buffer, 2.5 μl transposase (Tn5, 100 nM final), 22.5 μl water) was added to the nuclear pellet. Reaction was incubated at 37 °C for 45 minutes and amplified using PCR. Samples were purified using Qiagen MinElute PCR Purification Kit and again using Agencourt AMPure XP magnetic beads. For primer sequences, see **Supplementary Table 1**.

Samples were sequenced using Illumina NextSeq high output 2×75 bp settings. Seven samples were sequenced per run. Number of obtained sequencing reads is provided in **Supplementary Table 1**.

### ATAC-seq data quality control, alignment, and peak detection

Raw sequencing reads were inspected using fastqc version 0.11.7 (https://www.bioinformatics.babraham.ac.uk/projects/fastqc/) and subsequently trimmed with Trim Galore version 0.5.0 (https://github.com/FelixKrueger/TrimGalore) using parameters -- fastqc --paired --length 20 -q 20. Sequence alignment was performed using Bowtie2 version 2.3.4.1 (Langmead and Salzberg, 2012) against GRCh38 reference genome. During alignment, parameters --sensitive-local and -X 2000 were used. Additional filtering (-q 20), sorting and indexing was done with Samtools version 1.8 (Li et al., 2009). Finally duplicates were marked using Picard Markduplicates tool version 2.9.2 (http://broadinstitute.github.io/picard/), with parameters VALIDATION_STRINGENCY=LENIENT and REMOVE_DUPLICATES=FALSE. For filtered alignments, peak calling was done with MACS2 v2.1.0 (Zhang et al., 2008) using parameters -g hs --llocal 160000 --slocal 147 -q 0.05 -f BAMPE --nomodel --broad --bgd --call-summits. Final quality control was performed for aligned samples after peak calling using ataqv toolkit (version 1.0.0, https://github.com/ParkerLab/ataqv).

### Identification of artefact regions

As significant number of ATAC-seq reads originate from mitochondria, this can bias analysis at loci which have homology to autosomal or sex chromosome sequences. To exclude these regions from the analysis, we generated 100 copies of all the 30-mer sequences from mitochondrial DNA and aligned them to CRGh38 genome reference from which mitochondrial DNA had been excluded. Bowtie2 with --very-sensitive parameter was used. Alignments were converted to bed ranges using bedtools version 2.27.1 genomecov and merge tools (Quinlan and Hall, 2010).

### ATAC-seq signal quantification

We binned the genome into overlapping windows of size 500 bp and steps of 250 bp. To obtain read counts in each window, we used *bedtools coverage-counts*. For robust quantification of the signal in loci of interest, background correction, normalization and bias correction steps were performed. To obtain background corrected read count *c* for a given window at position *x*, we used the following formula:

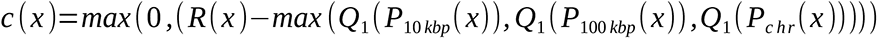

where *R* is the read count for the window at position *x, P_10kbp_()*, *P_100kbp_()*, and *P_chr_()* are lists of read counts for all windows within the range of +/-5kbp, +/-50kbp, and chromosome arm, respectively, from position *x*, excluding the window at position *x. Q_1_* is the value corresponding to the first quartile. This correction compensates for the variation in local background between samples and also enables detection of DARs from copy number aberrated genome areas (**Supplementary Figure 12A**). After background correction, we applied the median of ratio normalization (Anders and Huber, 2010), where sites with geometric mean below 1 were excluded from the calculation of the ratios, to obtain normalized read counts.

To compensate for potential bias due to sample collection procedure (RP and TURP), we divided the samples in the two groups. In the TURP group, we randomly assigned 4 BPH and 4 CRPC samples and in the RP group 4 BPH and 4 PC samples to keep the group sizes fixed. For each window, we applied the two-sided Wilcoxon rank-sum test. Random assignment of samples and significance testing was repeated 100 times. If 5th percentile of p-value distribution for a given window was less than p=0.01, we calculated the difference between medians of all the TURP and RP samples normalized read counts and subtracted this difference from all the TURP samples normalized read counts (717930 sites were corrected i.e. 6.5% of all sites). Application of this correction to normalized read counts resulted in quantified ATAC-seq signal.

### Identification of the differentially accessible regions (DARs)

To identify DARs, we compared the samples from two different groups (BPH to PC or PC to CRPC). We calculated the log2-ratio of the median value of each group (eg. log2(median(PC) / median(BPH))), absolute median difference between two groups (e.g | median(PC) - median(BPH)|), and used the two-sided Wilcoxon rank-sum test of two groups. For each window, we checked whether all the following 3 criteria were satisfied: |log2-ratio| > 2; p-value < 0.01; absolute-median-difference > 14. These thresholds were derived based on false discovery rate (FDR) analysis and correspond to FDR 9.7% and 9.14% in BPH to PC and PC to CRPC comparisons, respectively. If the log2-ratio of a DAR was positive, we called it an opening DAR and if the log2-ratio of a DAR was negative, we called it a closing DAR.

### Copy number aberration analysis

Raw sequencing reads from the whole genome sequencing experiment (DNA-seq) were aligned to the GRCh38 reference genome using Burrows-Wheeler Aligner (BWA) version 0.7.17 (Li and Durbin, 2009). Duplicate reads were marked using SAMBLASTER version 0.1.22 (Faust and Hall, 2014). Alignments were converted to BAM format and sorted using Samtools. We used Segmentum (Afyounian et al., 2017) to perform copy number analysis for the samples for which we had whole genome sequencing data (i.e. 4 BPH, 15 PC, 7 CRPC samples). Copy numbers were called using pooled BPH samples as reference with the following parameters: read depth were extracted for windows of width 500 bp, *window_size=15*, *clogr_threshold=0.8*, *min_read=35*, *logr_merge=0.2*. We used the reported log2-ratios for each genomic segment from Segmentum’s result to infer the copy number of that segment. This data was used to confirm that quantified ATAC-signal was not confounded by copy number alterations (**Supplementary Figure 12A**).

### Identification of the differentially methylated regions (DMRs)

Methylated DNA immunoprecipitation (meDIP) sequencing data was aligned to GRCh38 using Bowtie2 (settings: --score-min L,0,-0.15.), alignments were converted to BAM format and sorted using Samtools. Duplicated reads were marked with Picard Markduplicates. Samtools was used to filter out the duplicate reads. Differentially methylated regions were identified as described above for DARs using meDIP samples for which we had ATAC-seq data available. In the median of ratio normalization step, sites with geometric mean below 2 were excluded from calculating the ratios. DMRs were called with criteria |log2-ratio| > 2; p-value < 0.01; absolute-median-difference > 10, corresponding to FDR 4.61% and 7.90% for BPH to PC and PC to CRPC comparisons, respectively. If the log2-ratio of a DMR was positive, we called it a hypermethylated DMR and if the log2-ratio of a DMR was negative, we called it a hypomethylated DMR.

### Compilation and quantification of the peak set

In order to compile a consensus set of peaks across all samples, we adapted the approach from (Corces et al., 2018). For each individual sample, we used the summits position of peaks called by MACS2 (Zhang et al., 2008) and extended them by +/-250 bp to acquire the raw peak set for that sample. Preliminary signal for each raw peak was obtained using the above presented ATAC-seq signal quantification. If a raw peak was overlapping several adjacent windows, the weighted average based on the amount of overlap between the peak and overlapping windows, was used. For each sample, if there were overlapping raw peaks, the raw peak with the highest preliminary signal was selected. To standardize the peak signals across samples, these were further scaled in each sample by the sum of the signals of all the peaks divided by 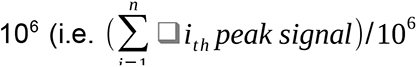 where *n* is the number of raw peaks in a given sample). Next, we pooled the peaks across all samples and removed their overlaps with the above approach using scaled signal values. Further, we removed raw peaks from the set if they were only present in one sample. This resulted in a peak set without overlaps.

To quantify the peaks signal, we used the approach above at the peak coordinates. A peak was removed from the peaks set, if all samples had standardized signals below a data-driven threshold (*t=5*) for that peak (**Supplementary Figure 12B**). Using this filtering criterion, we removed 4,935 loci. Finally, 127 peaks overlapping the artefact regions were removed. This resulted in a final 178,206 peak set for analysis.

### Quantification of chromatin accessibility at Transcription Start Sites (TSSs)

We extracted TSS coordinates for 18,537 protein coding genes and 1,471 miRNA (see quantification of gene and smallRNA expression below) from Ensembl version 90. For each gene and miRNA, we quantified chromatin accessibility within +/-500bp window from TSS using the same signal quantification approach as with the above peak set. The larger window size was used to account for the shape of the ATAC signal at the TSS sites (Supplementary Figure 1A).

### Visualization of the coverage at peaks, DARs and DMRs

All boxplots show the quantified ATAC-seq signal at peak or DAR locations. In all boxplots, the median is shown with a green line and mean with a red triangle. Lower and upper whiskers have been set to first quartile (Q1) - 1.5*IQR (interquartile range) and third quartile (Q3) + 1.5*IQR, respectively.

In coverage plots, we extended midpoints of loci of interest by +/-1.5 kbp. For the resulting regions in each sample, we extracted the read counts in bins of size 10bp using *bedtools coverage*. Next, we subtracted an estimated global background from the read count of each bin to acquire the background corrected read counts. To estimate the global background, we randomly selected 50,000 loci of size 500 bp excluding those that overlap with the loci of interest using *bedtools random*. We extended, binned and quantified each of these loci as above. The global background was calculated by the arithmetic mean across all the binned read counts from random loci. In case of meDIP data, if a bin had a background corrected read count above 50 across all samples, it was considered as an artefact region and the read counts for that locus were set to zero. To generate sample-specific profiles, we calculated the arithmetic mean of background corrected values across the corresponding bins for all loci of interest. Finally, we calculated the median across the corresponding bins of sample-specific profiles for each group (i.e. either BPH, PC, or CRPC).

### Annotation of peaks and DARs and DMRs

We annotated the loci of interest using *annotatePeaks.pl* routine from Hypergeometric Optimization of Motif EnRichment tool (HOMER; (Heinz et al., 2010)). We grouped regions annotated as 3’ UTR, TTS, non-coding, 5’ UTR, and exon under the term “Exon + untranslated”. We further annotated the loci of interest using *bedtools intersect* or *closest* with the following data sets. GeneHancer version 4.7 (Fishilevich et al., 2017) was used to annotate known regulatory elements (enhancers and promoters) and predicted regulatory region target gene associations. Pan-cancer peak set and PRAD peak calls from ATAC-seq data generated from TCGA samples (Corces et al., 2018), and Roadmap Epigenomics project DNase-seq data (Roadmap Epigenomics Consortium et al., 2015) were used to annotate previously identified accessible chromatin areas. Roadmap Epigenomics data was downloaded from reg2map (https://personal.broadinstitute.org/meuleman/reg2map/HoneyBadger_release/) and data from 3 distinct sets of regions (i.e. promoters, enhancers and dyadic regions) were combined. Duplicates were removed and LiftOver (Hinrichs et al., 2006) was used to convert the GRCh37 coordinates to GRCh38 (only 0.03% of the sites were lost due to LiftOver). To annotate ATAC features with experimentally validated transcription factor binding sites (TFBS), we downloaded the data from the Gene Transcription Regulation Database version 18.06 (Yevshin et al., 2019) which collects 5,068 ChIP-seq experiments and data from 846 unique TF. From the entire database, we subset GTRD ChIP-seq data for binding sites detected in prostate cancer cell lines and use it as a prostate-specific set, including 40 unique TFs from 1818 experiments. We assigned TFBS to ATAC features using R *findOverlaps* function.

We checked the overlap between DMRs and the CpG islands using the information obtained from UCSC genome browser (http://hgdownload.cse.ucsc.edu/goldenpath/hg38/database/cpgIslandExt.txt.gz) by using *bedtools closest*. If the distance of a locus and its closest CpG island was below 2 kbp, we marked that locus as a CpG island.

We retrieved a list of 3,804 human housekeeping genes from (Eisenberg and Levanon, 2013). We mapped gene names to Ensembl gene id version 90 using the R *merge* function. The resulting list included 3,662 genes.

### Quantification of gene expression

Previously published transcriptome sequencing data, including 12 BPH, 30 PC and 13 CRPC samples (Ylipää et al., 2015) was aligned to GRCh38 and quantified by STAR version 2.5.3a using Ensembl version 90 annotations. We obtained quantification of 58243 genes. Samples with high quality data were matched with ATAC-seq samples (**Supplementary Table 1**). Lower quartile value of expression distribution across all the samples was used as a threshold to remove low expressed genes, resulting in 18537 protein coding genes (mitochondrial excluded) available for the analysis. The DESeq2 version 1.20 Bioconductor package was used to model the data and extract differentially expressed genes. We fit the model taking into account both RNA isolation methods (Qiagen™ Trizol™ and Qiagen™ All Prep™) and stages of prostate cancer progression (BPH, PC, CRPC). To address the bias introduced by different extraction protocol, we used coefficients estimated from the model (Ylipää et al., 2015): we extracted coefficients for the RNA isolation method covariate using DESeq2 *coef* function and subtracted these values from library size corrected read counts of Trizol-treated samples in log2 scale. We detected differential expression in two comparisons, BPH to PC and PC to CRPC. A gene was considered differentially expressed (D.E.) if the absolute median difference of normalized read counts between the groups was greater than 180, the log2 fold change was greater than 1 and the FDR corrected p-value lower than 0.05. In the BPH to PC comparison and PC to CRPC comparisons 933 and 533 D.E. genes were detected. If the log2-ratio of a D.E. was positive, we called it an overexpressed D.E. gene. If the log2-ratio of a D.E. was negative, we called it an underexpressed D.E. gene.

### Quantification of small RNA expression

Previously published small RNA sequencing data (Ylipää et al., 2015) was re-analysed by mapping sequence tags to human sequences from mirBase version 22. We mapped sequencing tags allowing for single base deletion at the 3’ or insertion at either 3’ or 5’. Modified sequences mapping to the same mirBase identifier were collapsed and their abundance summed. This process yielded data for 1471 annotated miRNA sequences. The resulting data matrix was normalized using median of ratios normalization, genes with geometric mean lower than 15 were discarded. Differentially expressed miRNA were detected in BPH to PC and PC to CRPC comparisons. A miRNA was considered differentially expressed if showing a log2 fold change greater than 1 and a FDR adjusted t-test p-value lower than 0.05. This analysis yielded 26 and 51 differentially expressed miRNA for BPH to PC and PC to CRPC comparisons, respectively.

### Protein expression data

We used our previously published sequential window acquisition of all theoretical mass spectra (SWATH-MS) data and defined differentially expressed proteins as described earlier (Latonen et al., 2018).

### Quantification of AR activity score

AR activity score was determined using a publicly available gene expression signature composed of 27 genes (Hieronymus et al., 2006). Of these, 21 genes are upregulated in the first 24 hours after androgenic treatment: PSA, TMPRSS2, NKX3-1, KLK2, GNMT, TMEPAI, MPHOS9, ZBTB10, EAF2, BM039, SARG, ACSL3, PTGER4, ABCC4, NNMT, ADAM7, FKBP5, ELL2, MED28, HERC3, MAF.

Normalized gene expression values in log2 scale were converted to z-scores:

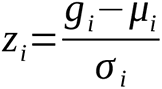

where *i* represents a gene from the list above, *g* the gene expression, *μ* the arithmetic mean of expression values and *σ* the standard deviation of the gene. Both mean and standard deviation were computed using all samples. For each sample, AR activity score was computed by summing genes scores:

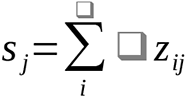

Scores were split according to sample groups: BPH, PC and CRPC. Score distributions were visualized as violin plots. Both upper and lower quartiles and the mean activity score value were overlaid on the figures. P-values were computed with two tails Mann-Whitney U-test to assess the statistical significance between BPH and PC or PC and CRPC scores under the null hypothesis of no difference between groups.

### Association of chromatin accessibility with target genes

To link chromatin features (peaks and DARs) to putative target genes, we performed correlation analysis across all samples with RNA-seq, smallRNA-seq, or SWATH-MS protein data. We calculated Pearson and Spearman correlations between ATAC-seq signal and gene or protein expression in four different contexts: 1) at transcription start site (TSS), defined as +/-500bp from TSS annotation, we computed correlation between TSS ATAC-seq signal and corresponding gene expression; 2) we defined a region of 1 kbp upstream and 100 bp downstream of TSS as a promoter, we searched for ATAC features overlapping this region and computed correlation between their signal and corresponding gene or protein expression, if available; 3) for each ATAC feature we searched for the closest gene using annotations from the HOMER tool and computed correlation between their signal and gene or protein expression, if available; 4) we used all ATAC features and genes falling within same TAD to compute correlation between all pairs. To define TAD boundaries, we used annotations from ENCODE consortium based on data from LNCaP cell line (ENCODE Project Consortium, 2012), GEO accession: GSE105557, downloaded from http://promoter.bx.psu.edu/hi-c/ (Wang et al., 2018)). We extended this list with genomic intervals included between each pair of TAD using *bedtools complement* and merged the resulting list with the initial one.

To derive a threshold for significant associations, in each context, we computed null distributions by randomizing sample order prior to correlation coefficient computation (**Supplementary Figure 9A**). To enable false positive rate estimation, randomization was repeated 10 times for each pair of comparisons in each context. Based on evaluation of the distributions, we chose to set thresholds to |correlation coefficient| > 0.5 for genes and | correlation coefficient| > 0.6 for proteins resulting in false positive rate from 5.4 * 10^-3^ to 1.3 * 10^-3^, respectively (**Supplementary Figure 9A**). Associations with either Pearson or Spearman correlation above threshold were kept for the downstream analysis. The above analysis was implemented by custom script using standard Unix tools, Python 3.6.8, R version 3.5.2 and packages from the Bioconductor framework managed via the BiocManager package version 3.8, HOMER tool and *bedtools*.

### TF gene expression network

Each gene was assigned to one or more ATAC-seq features from previous correlation analysis. Transcription factor binding sites in ATAC-seq features were detected during the annotation step. A gene was defined as the target of a transcription factor if its expression showed correlation with accessibility of an ATAC-seq feature carrying a binding site for the TF. For each pair of TFs, the number of co-regulated genes was calculated resulting in a contingency matrix of 845×845 TFs. This matrix was filtered to retain TFs sharing at least 100 genes, leaving a 192×192 contingency matrix. Hierarchical clustering was applied and two clusters were detected. Manhattan distance was used as distance metric and UPGMA as clustering algorithm (**Supplementary Figure 10A**). The smallest cluster, containing the majority of detected connections, was extracted. Another filter was implemented similarly to the one described above: TFs sharing at least 300 genes with at least one other factor were retained. This filtering procedure resulted in a 41×41 contingency matrix. For visualization hierarchical clustering was calculated using Euclidean distance as distance metric and complete linkage as clustering algorithm. The analysis was implemented in R 3.5.2 using the *pheatmap* package for visualization and basic clustering functions.

### TF binding site enrichment analysis

We determined the number of clusters for k-means clustering using consensus clustering with elbow method. For clustering, we used top 20% peaks with highest variance. For relative TF enrichment analysis, each cluster was compared against all the others. Enrichment analysis was performed using HOMER *findMotifs.pl* version 4.10. We used the full HOCOMOCO version 11 human TF (p 0.001) (Kulakovskiy et al., 2018) database as a known input TF list. Plotting was done using the R 3.5.2 and ggplot package.

### TF footprinting and accessibility

For TF footprint depth and flanking accessibility analysis Tn5 cut sites were counted using custom R scripts. Pooled samples for BPH, PC and CRPC groups were generated using *Picard MergeSamFiles* and used for group level analysis. In other analysis, individual BAM files were used directly. Possible TF binding locations were predicted using FIMO version 5.0.2 (Grant et al., 2011) with HOCOMOCO v11 database and *--thresh 0.001* parameter. Predicted sites were intersected with peaks and DARs from BPH to PC and PC to CRPC comparison groups. We filtered the TF list by gene expression across samples. TF belonging to the lower quartile of this distribution were discarded. We quantified footprint base as mean count of insertions at the motif positions, while for flanking height, we considered 25 bases around each detected motif. To quantify each motif background, we used a set of 25 bp windows 200 bp upstream and downstream of the motif center. We computed flanking accessibility as log2(flanking height/background) and footprint depth as log2(footprint base/flanking height). For expression association, Pearson correlation between these footprint parameters and TF expression was calculated. In footprint visualization, the number of cutting sites were scaled according to read numbers in respective phenotypes.

### Motifs discovery from accessible chromatin sites

We used the BPNET Python package version 0.0.21 (Avsec et al.) to train and interpret sequence-to-profile convolutional neural networks from sample-specific ATAC-seq data. In BPNET recurring patterns with high contribution scores are clustered based on sequence identity to build contribution weight matrices (CWMs). We first tested the applicability of the BPNET model with data from VCaP cell lines. We compared the CWMs obtained from models trained with publicly available AR Chip-seq data (Massie et al., 2011) and with ATAC-seq data generated in-house. These data were aligned and peaks detected as presented above. To consider the TF-specific binding context in ATAC-seq data, we extended the ATAC-seq peaks summits by 50 bp in both directions, and intersected the 100 bp regions with the direct AR-DNA interaction map defined in the UniBind database (Gheorghe et al., 2019). We kept the regions having an intersection of at least 1 bp, and selected these peaks as model training sequences. We tested the similarity between the resulting CWMs and all the known Position Weight Matrices (PWMs) collected in the HOCOMOCO v11 database (Kulakovskiy et al., 2018) using the Tomtom motif comparison tool (Gupta et al., 2007). Tomtom results were used to identify the TF or TF family to be associated with each CWM. We observed that the motifs discovered by the model trained with Chip-seq data were also discovered by the model trained with ATAC-seq data (**Supplementary Figure 4**). We trained BPNET models for each of the 38 clinical samples using the above presented peaks set summits to define the training sequences and the ATAC-seq data. We then applied the procedure we tested on cell lines to build and interpret TF-specific models for 4 TFs - namely AR, FOXA1, HOXB13, and ERG - on the highest quality samples (6 BPH, 4 CRPC, 8 PC) having TSS enrichment > 3.5. ATAC-seq BPNET models were trained on canonical chromosomes with default hyper-parameters, and chromosomes 2, 3, and 4 as validation chromosomes. Chip-seq BPNET models were trained using the same configuration, except an increased kernel size of 50 for the transposed convolution layer. The models trained in cell lines and the models trained in clinical samples using the above presented peaks set summits were trained with a patience of 5 epochs. The TF-specific models trained in clinical samples were trained with a patience of 20 epochs. We represented the information content of the discovered motifs as sequence logos using the built-in BPNET function; when more than one meta cluster was reported by BPNET, we omitted the meta clusters with no matching TFs if at least one pattern in another meta cluster had a TF or a TF family assigned to it (**Supplementary Figures S5-S8**).

## Data and code availability

Sequencing data has been deposited in European Genome-phenome Archive under accession number EGAS00001000526. Code used for the analysis is available at https://github.com/nykterlab/Tampere_PC/

## Authors contributions

Initiated the study: MN

Supervised the work: MN, TV, KG, JK

Designed the analysis: MN, KG, JK, LL, AU

Produced the data: JU

Contributed data / samples / analysis tools: MA, KK, TT

Performed analysis: JU, EA, FT, TH, AL, AS, KG, RN

Drafted the manuscript: JU

Wrote the paper: JU, EA, FT, TH, KG, MN

All authors have read and approved the final version of the paper.

## Declaration of interests

All authors declare that they have no conflicts of interest.

## Acknowledgements

We thank Paula Kosonen, Hanna Selin, Riina Kylätie, Marja Pirinen, and Päivi Martikainen for their technical assistance. Financial support is acknowledged from: Academy of Finland project no. 312043 (MN), project no. 310829 (MN), project no. 333545 (KG), project no. 317871 (LL), Cancer Foundation Finland (MN, KG, LL), Competitive State Research Financing of the Expert Responsibility area of Tampere University Hospital (MN, KG), Norwegian Cancer Society project no. 198016-2018 (AU), Pirkanmaa Regional Fund (JU), Tampere University doctoral school (EA, FT), Emil Aaltonen Foundation (KG), Sigrid Juselius Foundation (MN, KG, LL), Foundation of the Finnish Cancer Institute (LL). CSC—IT Center for Science Ltd. (https://www.csc.fi/csc) provided computational resources for the work. The results published here are in part based upon data generated by The Cancer Genome Atlas project (dbGaP Study Accession: phs000178.v9.p8) established by the NCI and NHGRI. Information about TCGA can be found at http://cancergenome.nih.gov. We acknowledge ENCODE Consortium and the ENCODE production laboratories for generating the Hi-C data.

## Supplementary Figures

**Figure S1.**
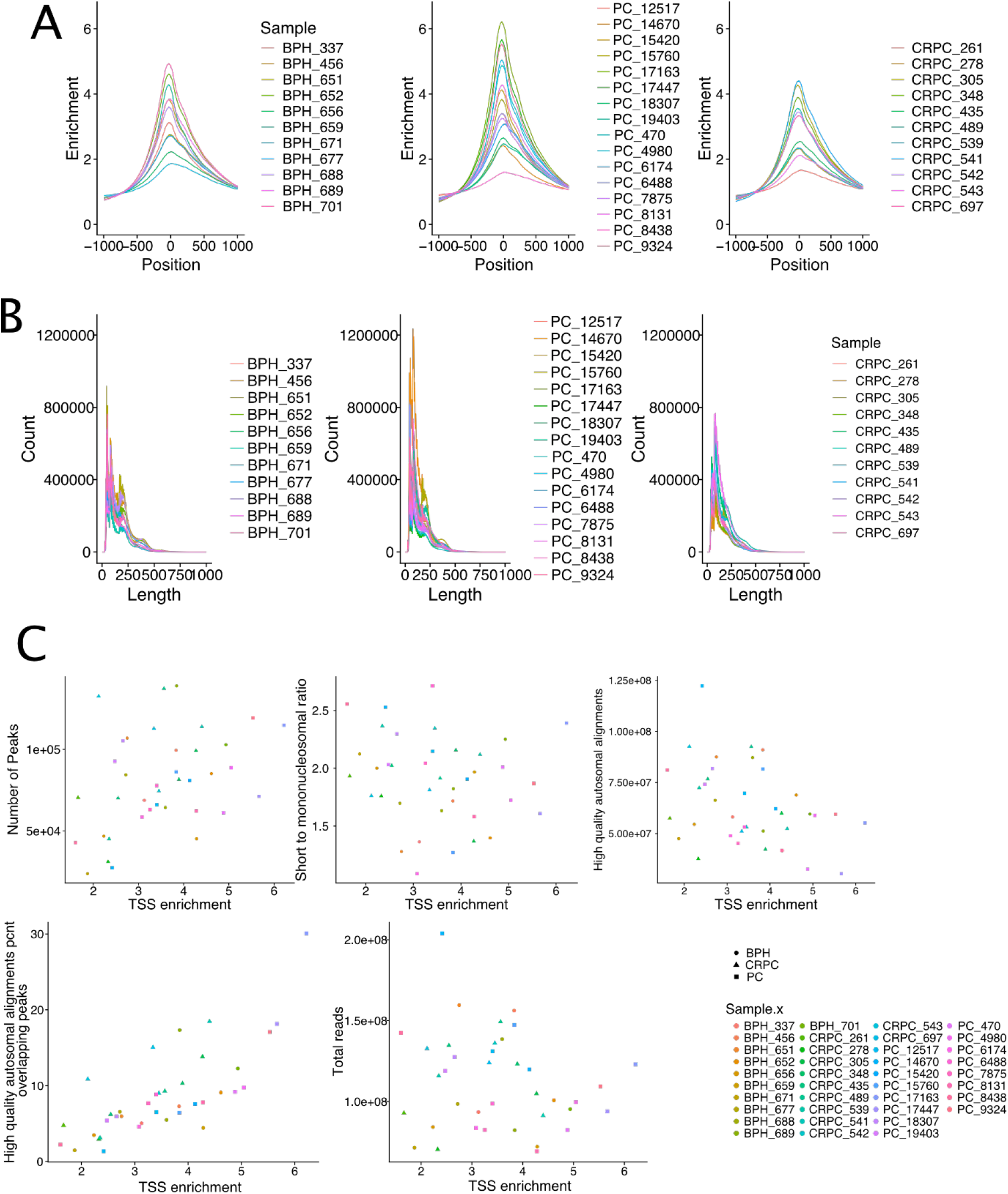
**A.** ATAC-seq enrichments at TSS for all samples. Sample groups have comparable quality and variability. **B.** Fragment length patterns in all samples show the first peak around 160 bp, matching single nucleosome size. A second peak is observed for the second nucleosome. **C.** Correlations between TSS enrichment and several key quality control values. The “high-quality autosomal alignments percentage overlapping peaks” value correlates with TSS enrichments, as expected. These show that no bias was introduced by the sequencing step.

**Figure S2.**
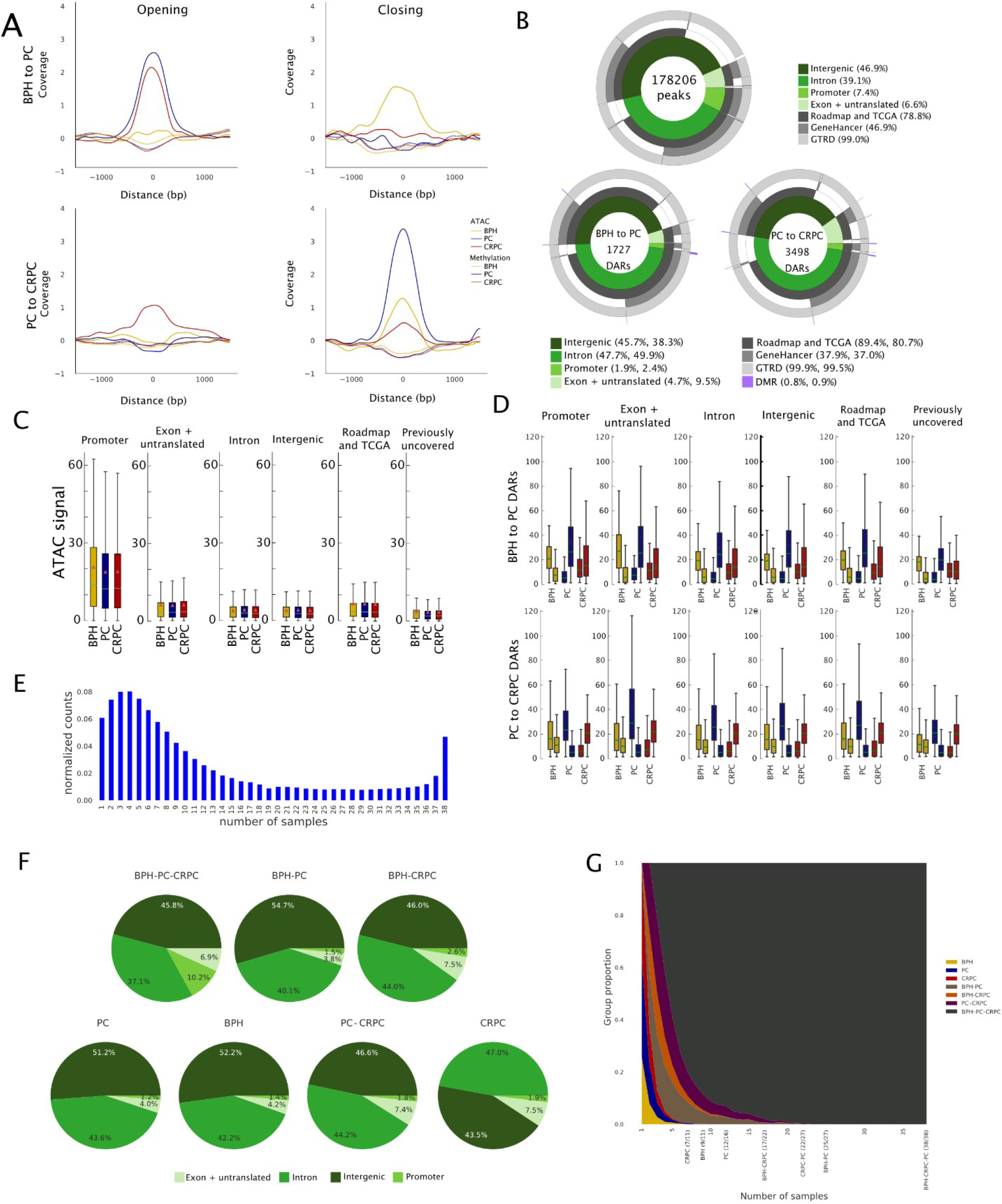
**A.** Background-corrected ATAC-seq and MeDIP coverage for loci corresponding to opening (left column) and closing (right column) DARs for BPH to PC (top row) and PC to CRPC (bottom row) comparisons. Each curve’s baseline has been shifted to zero for presentation clarity. The curves have been smoothed with a Gaussian filter with a standard deviation set to 7. **B.** Donut plots for locations of peaks and DARs from both comparisons. Majority of peaks and DARs are located in intergenic and intronic regions but there is a clear difference in promoter regions where ∼7.5% of peaks are located compared to ∼2% in both DAR comparisons. Overlaps with DMR regions are shown in the DAR donut plots. **C.** Peaks ATAC-signal in different annotation categories. Strongest ATAC-seq signal is detected at the promoters. **D.** Comparable ATAC-seq signal across different genome annotation areas. For each sample group, the left boxplot shows the signal in closing and right boxplot in opening DARs from respective comparisons. Data from the group of samples that were not part of the comparison (CRPC in top panels, BPH in lower panels) are shown from the same loci for reference. **E.** Normalized peak counts present in different numbers of samples. **F.** Genomic locations of peaks belonging to each sample group or combination of sample groups. Peaks belonging to the set with all sample groups have over 10% of peaks annotated to promoters, whereas in other groups the promoter fraction is 1.2-2.6%. **G.** Number of samples reporting peaks by group or combination of groups membership. The number of samples in which a peak is present is shown on the X-axis. Labels for sample groups and sample group combinations are reported. In addition, points where the number of peaks in groups or group combinations go to zero are also shown. The proportion of peaks in each group is shown on the Y-axis.

**Figure S3.**
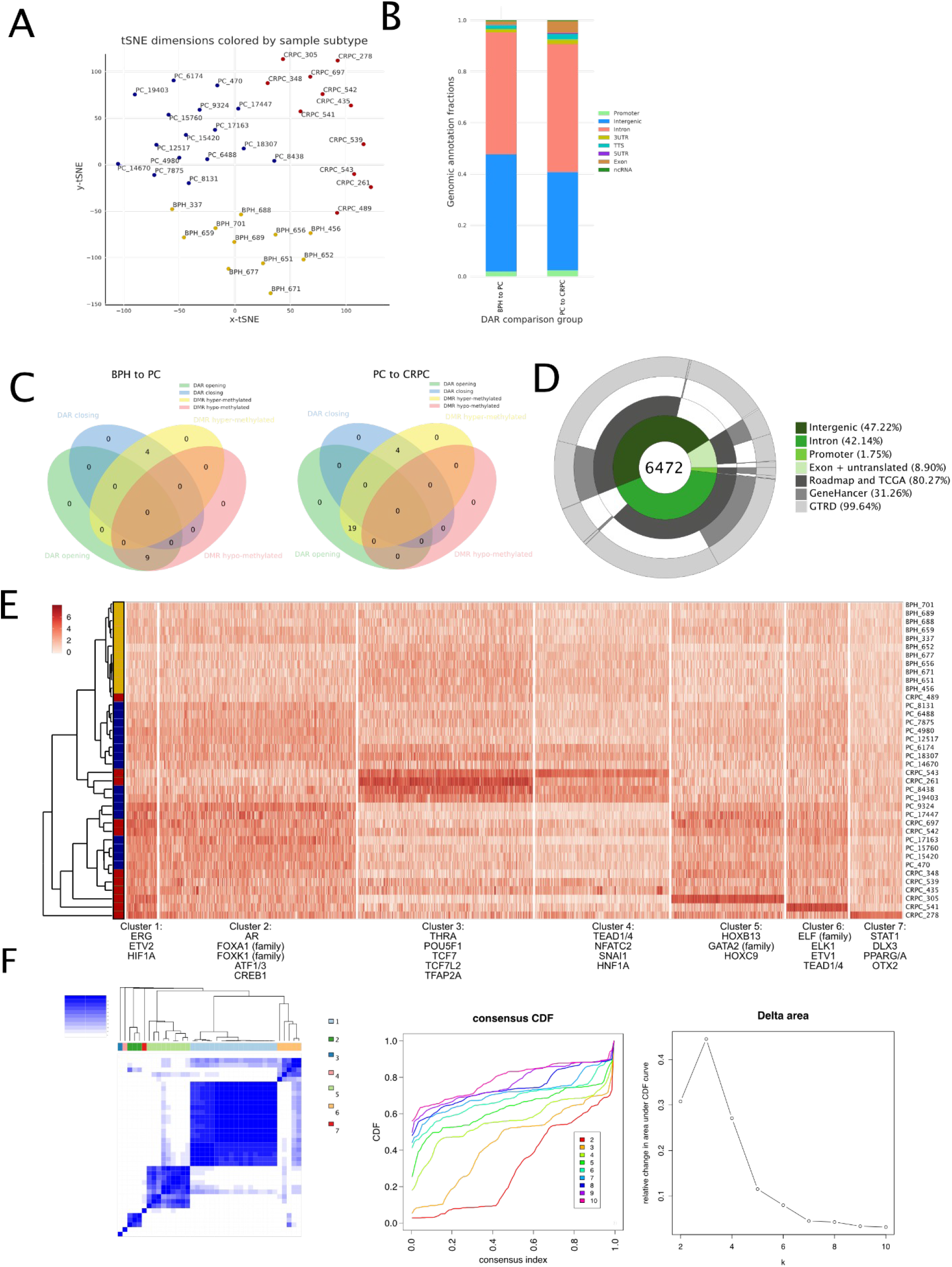
**A.** Two-dimensional t-SNE dimension reduction using normalized ATAC-seq signal from all DARs separates samples to their respective groups. Python’s Scikit-Learn package t-SNE algorithm implementation was used with default parameter values except *perplexity=15, metric = Pearson correlation,* and *method = exact*. **B.** Annotation of different genomic locations for DARs detected in BPH to PC and PC to CRPC comparisons. A higher fraction of DARs are located in the gene body and near the gene body in PC to CRPC compared to BPH to PC. **C.** Number of overlaps between DARs and DMRs in BPH to PC and PC to CRPC comparisons show only minimal overlap. **D.** Donut plot showing genomic annotations of cancer-specific peaks and overlap with previously reported features. **E.** K-means consensus clustering of the 20% topmost peaks with the highest variance identifies 7 clusters. Scale bar indicates quantification value. Examples of disease-relevant TFs from TF binding site enrichment analysis are shown for each cluster. **F.** Selection criteria for K=7 clusters in the cluster analysis. Consensus matrix (K=7, left), cumulative distribution function (CDF) plots for K=2-10 (middle), and relative change in CDF (right) are shown. K=7 illustrates stable cluster structure with relative CDF change at elbow point of the curve.

**Figure S4.**
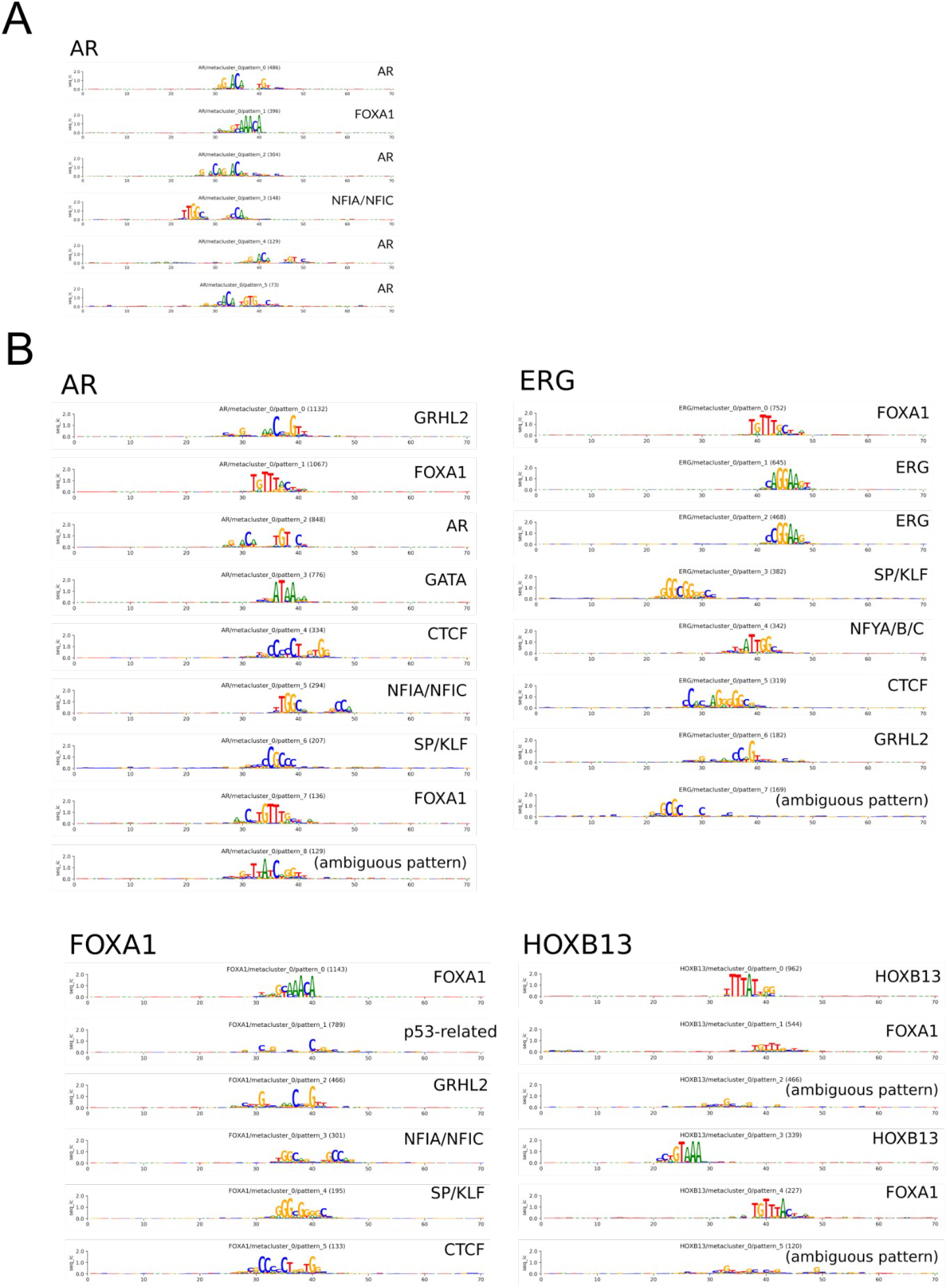
**A.** Binding motifs discovered with BPNET from AR ChIP-seq data generated from VCaP cell line. **B.** Discovered binding sites using ATAC-seq data from VCaP cells with binding sites for AR, FOXA1, HOXB13, and ERG.

**Figure S5.**
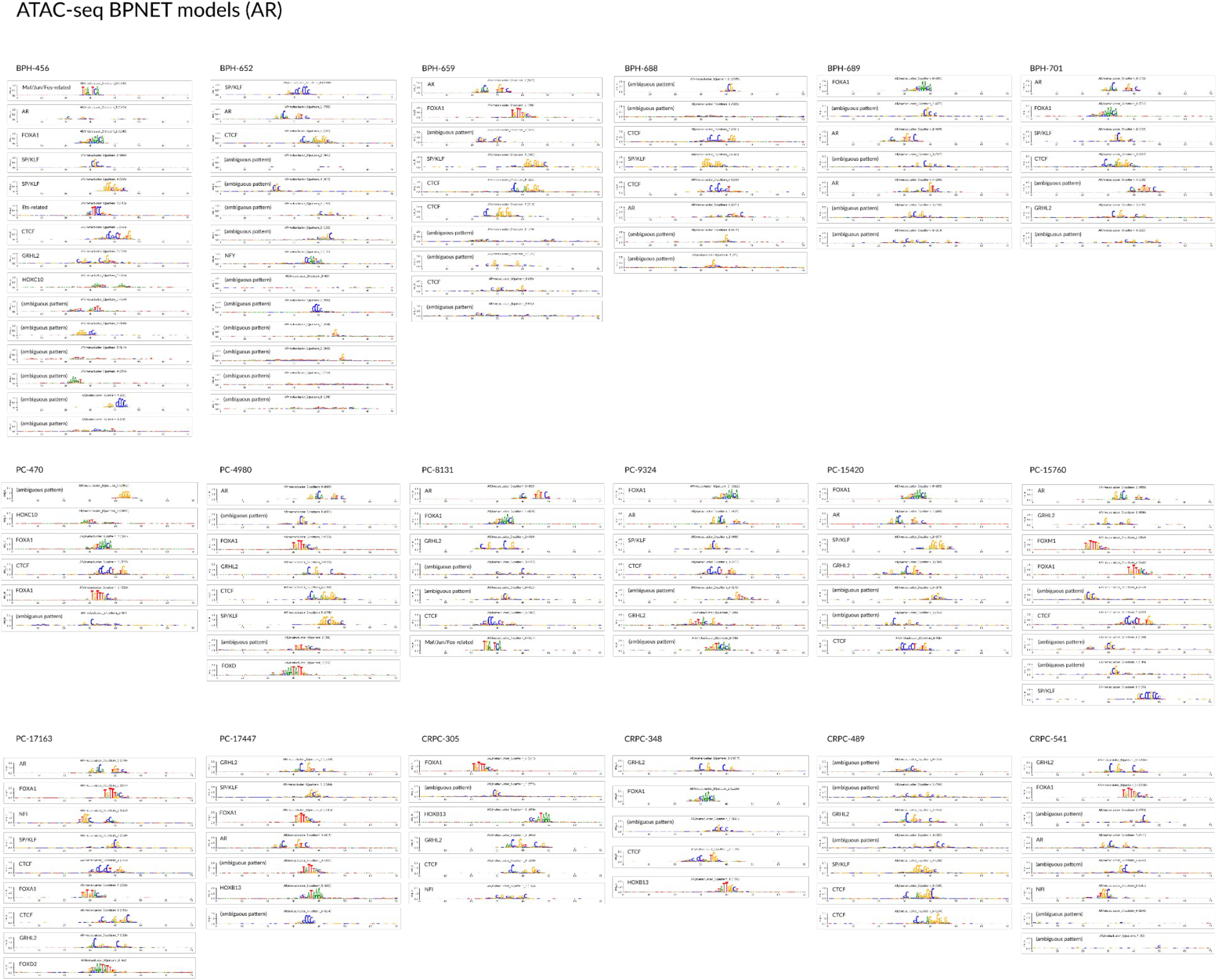
Binding motifs discovered with BPNET from clinical ATAC-seq samples using AR binding sites overlapping with peaks from ATAC-seq data. For each pattern, the number of BPNET *seqlets* contributing to that pattern is reported in parenthesis (see **Methods**).

**Figure S6.**
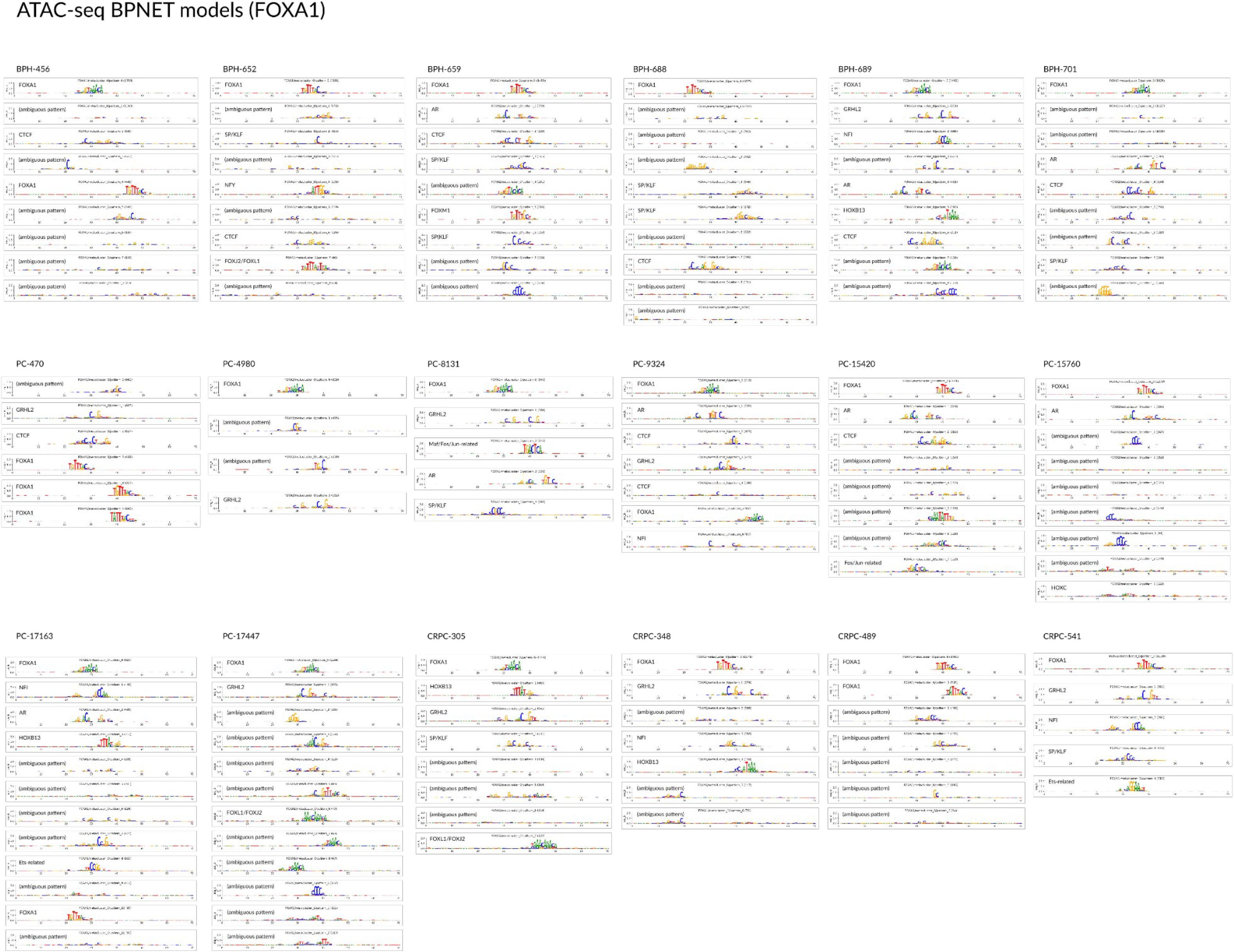
Binding motifs discovered with BPNET from clinical ATAC-seq samples using FOXA1 binding sites overlapping with peaks from ATAC-seq data. For each pattern, the number of BPNET *seqlets* contributing to that pattern is reported in parenthesis (see **Methods**).

**Figure S7.**
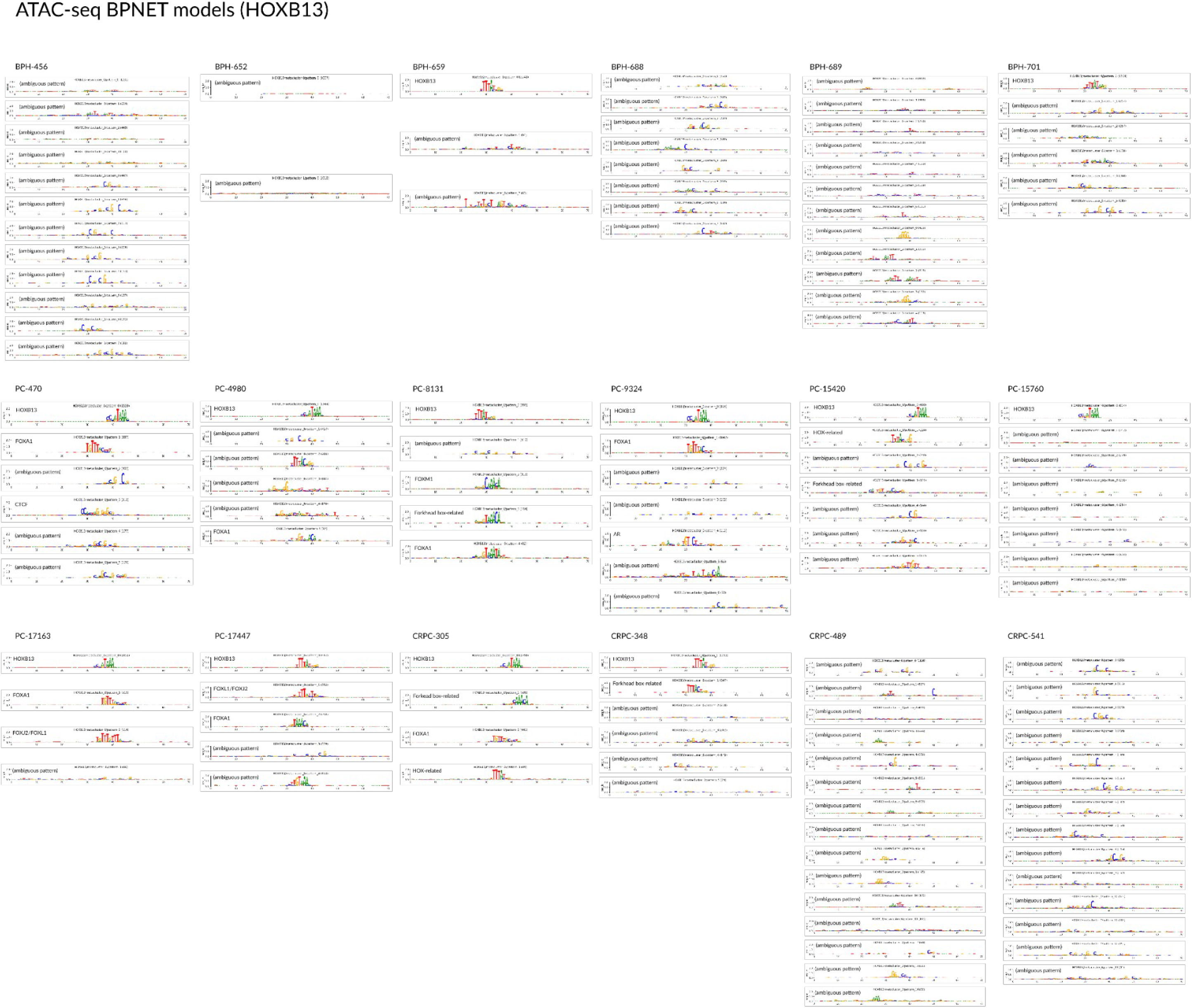
Binding motifs discovered with BPNET from clinical ATAC-seq samples using HOXB13 binding sites overlapping with peaks from ATAC-seq data. For each pattern, the number of BPNET *seqlets* contributing to that pattern is reported in parenthesis (see **Methods**).

**Figure S8.**
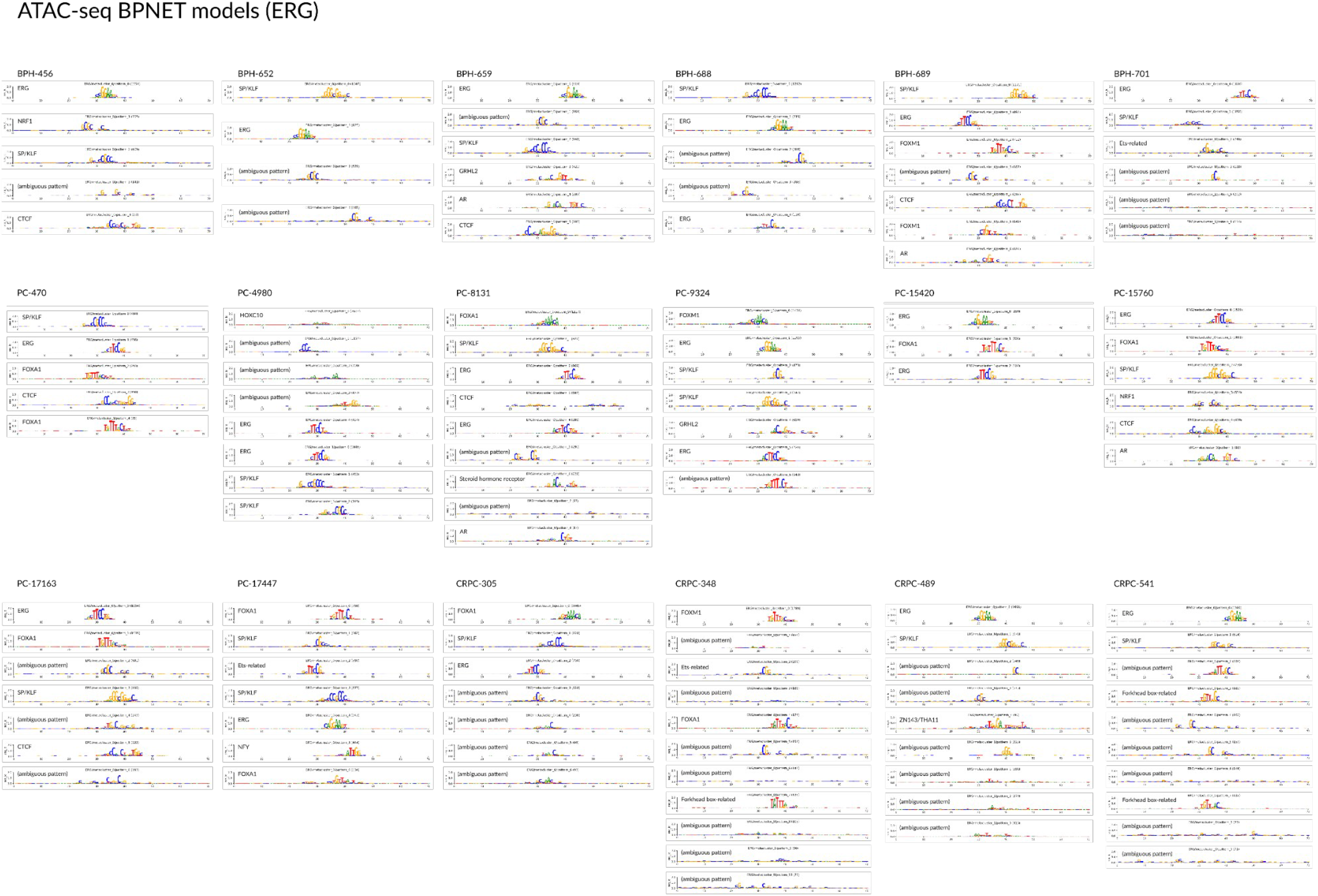
Binding motifs discovered with BPNET from clinical ATAC-seq samples using ERG binding sites overlapping with peaks from ATAC-seq data. For each pattern, the number of BPNET *seqlets* contributing to that pattern is reported in parenthesis (see **Methods**).

**Figure S9.**
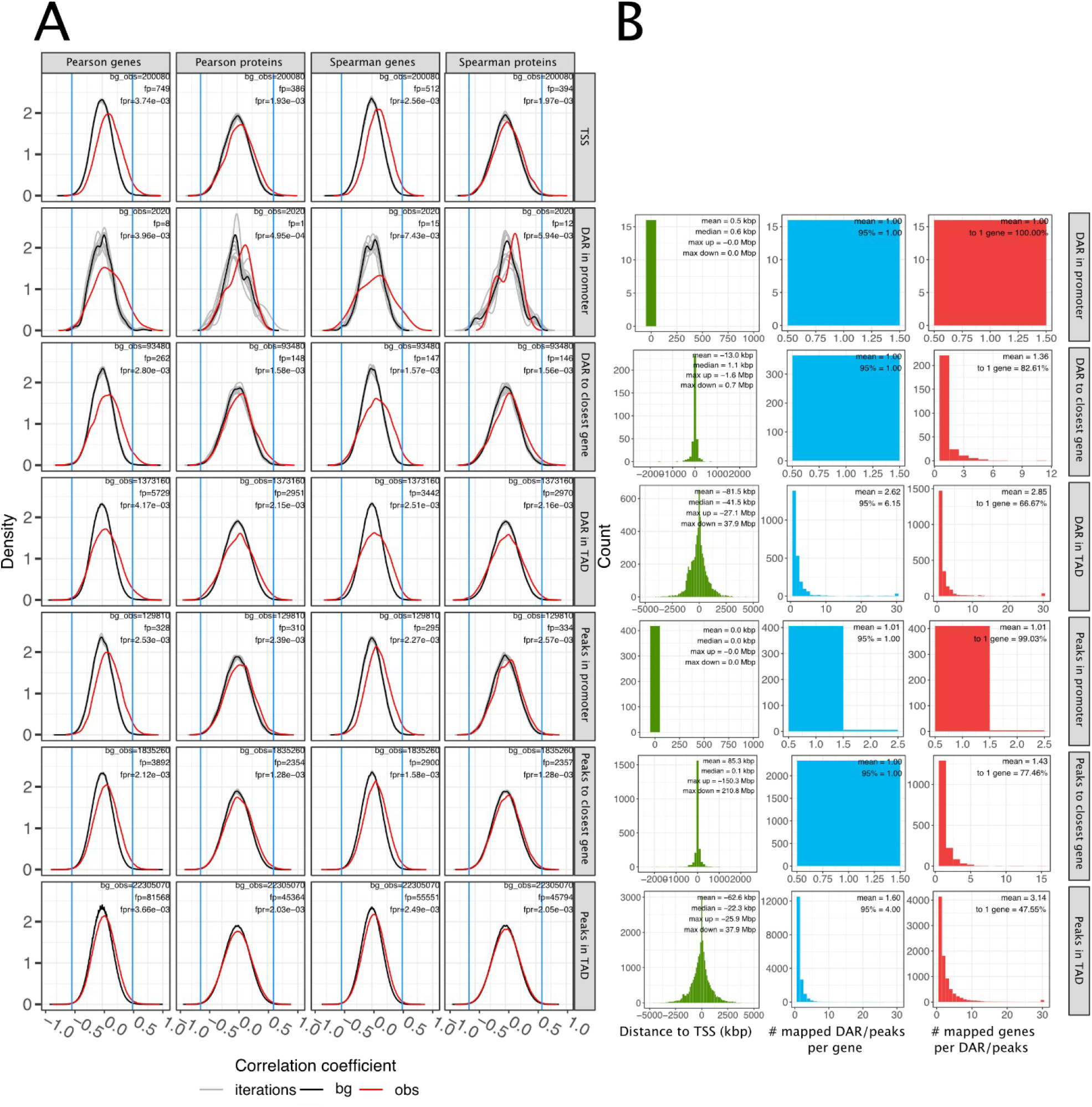
**A**. Pearson and Spearman correlation coefficients for all associations between chromatin accessibility and gene or protein expression. Data are shown for both gene (RNA-seq and smallRNA-seq) and protein expressions (SWATH-MS). Number of correlation coefficients used for null distribution, false positives and false positive ratio are reported in the inset. **B**. Histograms of distances between ATAC-seq features and TSS of the associated gene are shown across all comparisons. In addition, numbers of peaks/DARs associated to a given gene and also the number of genes associated to a given peak/DAR are shown as histograms (truncated at 30). Mean of these histograms is given in the figure. In addition, 95th percentile and fraction of peaks/DARs linked to a single gene are given in respective comparisons. Mean, median and maximum upstream (max up) and downstream (max down) distances are reported for the distance distribution. The percentage of ATAC-features linked to exactly one gene is also reported for the right panel.

**Figure S10.**
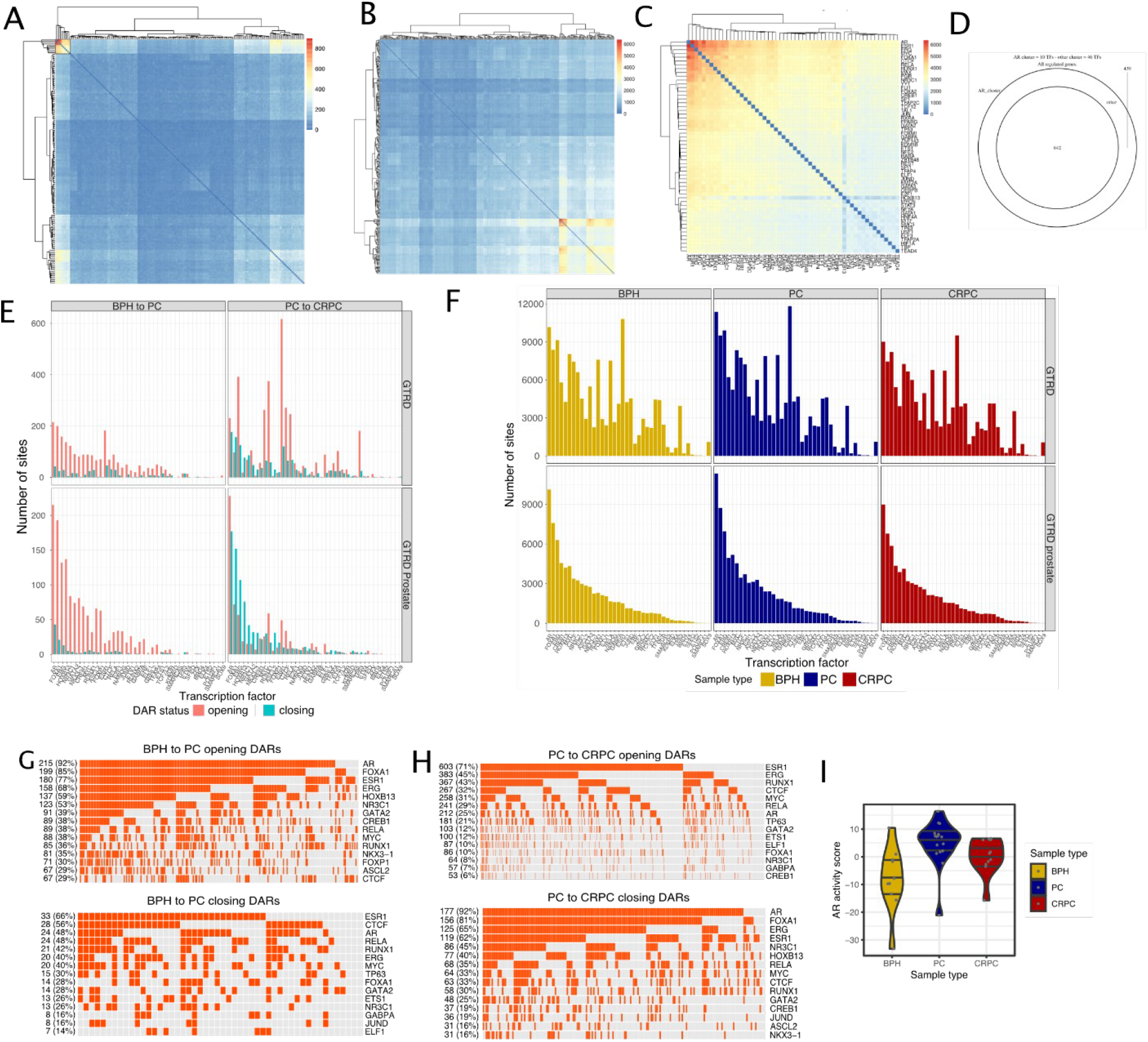
**A.** The heatmap represents a gene expression regulatory network created by using DARs and genes that have correlation: rows and columns are transcription factors (nodes), each cell in the matrix represents an edge, the weight of the edge is given by the number of shared genes which is encoded in the color. Rows and columns are filtered to have at least one cell with a value greater than 100. **B.** Same as panel A but using peaks. **C.** Subset of TFs with the highest number of genes (from data shown in panel B). The highest number of genes can be seen in the top right corner where there are the same four TFs as in Figure 4B. **D.** AR cluster-regulated genes from C are a superset of the genes regulated by the other cluster of TFs. The Venn diagram reports the agreement between the sets of genes regulated by the two clusters. **E.** Number of TFs that have binding site in DARs with associated target genes. Data are shown using all the data from GTRD (top panels) and using only prostate cancer-specific subset (GTRD prostate; bottom panels). Shown are both BPH to PC (left panels) and PC to CRPC (right panels) comparisons. **F.** Number of TFs that have binding site in peaks with associated target genes. Data are shown using all the data from GTRD (top panels) and using only prostate cancer-specific subset (GTRD prostate; bottom panels). In the GTRD prostate, we see that several key TFs like AR, FOXA1, ERG and HOXB13 are among the most common ones. **G.** Oncoprints representing TF binding sites overlapping DARs correlated with gene expression in BPH to PC comparison using the complete GTRD dataset. Number of binding sites for each TF is shown. In parentheses, the percentage of DARs reporting that binding site is also shown. **H.** Same as panel G but for PC to CRPC comparison. **I.** Violin plots of AR activity scores for each sample group. Individual samples are shown as grey dots.

**Figure S11.**
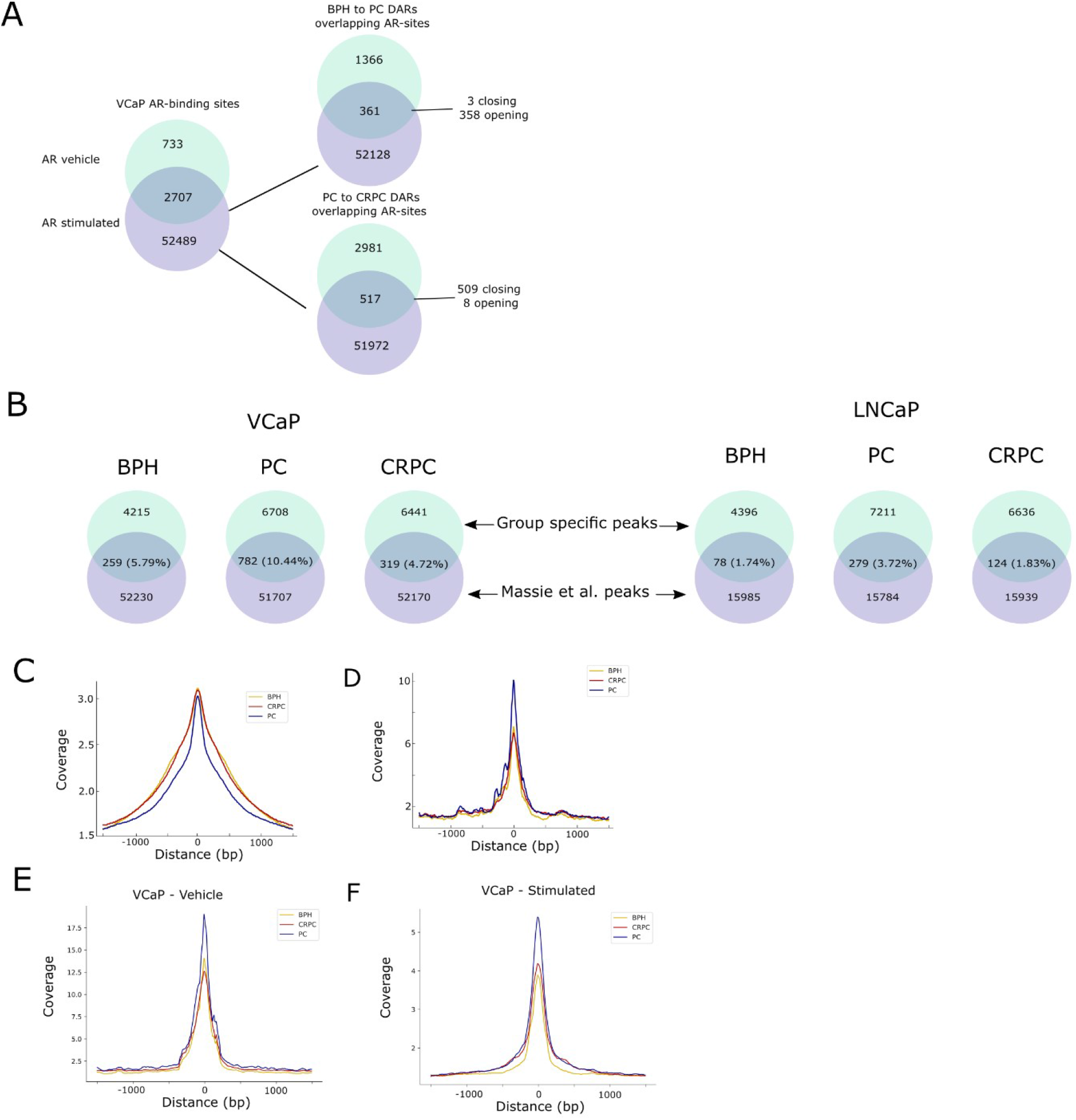
**A.** Number of sites and their overlap from DHT- and vehicle-treated VCaP cells. Sites that are present in DHT-stimulated cells are compared to DARs. In BPH to PC, most of the sites overlap with opening DARs and in PC to CRPC with closing DARs. **B.** Comparison of BPH, PC, and CRPC group-specific peaks to AR-stimulated peaks from LNCaP and VCaP cell lines <u>(Massie et al., 2011)</u> shows the highest overlap with the PC group. **C.** Background-corrected ATAC-seq coverage of AR binding sites from all GTRD AR binding sites. **D.** Background-corrected ATAC-seq coverage of AR binding sites from vehicle-treated cells (Massie et al. 2011). **E.** Background-corrected ATAC-seq coverage at AR binding sites from vehicle-treated VCaP cells. **F.** Same as panel E but with DHT stimulation. Signal is stronger in PC samples than other sample groups.

**Figure S12.**
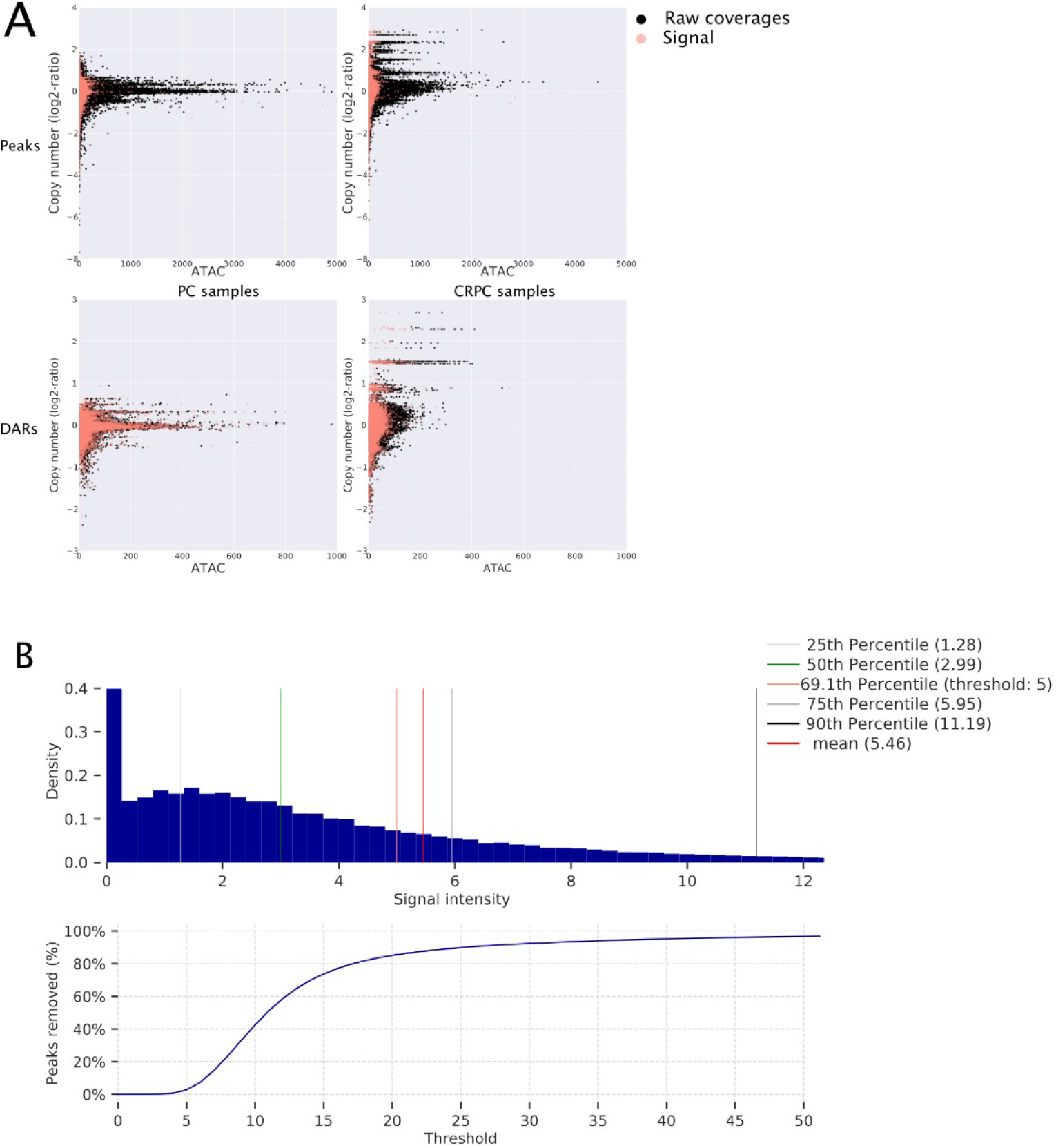
**A.** Effect of the background correction and normalization (Signal, red dots) in relation to raw ATAC data (black dots) and DNA copy number. Background correction and normalization successfully removes the linear relationship between copy number and ATAC coverage. **B**. Distribution of peak quantifications across samples. Different percentiles and the utilized threshold 5 are shown (top). The number of sites that would be removed with a given threshold (bottom).

## Supplementary Table legends

**Supplementary Table 1: Quality control metrics and peak detection results**

Table contains information about samples, and relevant information from sequencing such as quality control metrics and primers used. In addition, it contains information about peaks and their clustering.

**Supplementary Table 2: Differentially accessible and differentially methylated regions**

Table contains information about differentially accessible and differentially methylated regions in different comparison groups.

**Supplementary Table 3: TF binding analysis**

Table contains information about transcription factor footprint analysis, correlation between footprint depth, flanking accessibility and gene expression as well as information about motifs discovered using BPNET.

**Supplementary Table 4: Correlations of accessible chromatin regions and gene expression**

Table contains information about peaks and DARs correlation coefficients computed against gene and protein expression in different biological contexts. Table also reports gene names of transcription factors with binding site overlapping peaks and DARs and basic annotations of correlated genes.

## Notes

### Competing Interest Statement

The authors have declared no competing interest.

## References

Adams, E.J., Karthaus, W.R., Hoover, E., Liu, D., Gruet, A., Zhang, Z., Cho, H., DiLoreto, R., Chhangawala, S., Liu, Y., et al. (2019). FOXA1 mutations alter pioneering activity, differentiation and prostate cancer phenotypes. Nature 571, 408–412.

Afyounian, E., Annala, M., and Nykter, M. (2017). Segmentum: a tool for copy number analysis of cancer genomes. BMC Bioinformatics 18, 215.

Anders, S., and Huber, W. (2010). Differential expression analysis for sequence count data. Genome Biol. 11, R106.

Annala, M., Kivinummi, K., Tuominen, J., Karakurt, S., Granberg, K., Latonen, L., Ylipää, A., Sjöblom, L., Ruusuvuori, P., Saramäki, O., et al. (2015). Recurrent SKIL-activating rearrangements in ETS-negative prostate cancer. Oncotarget 6, 6235–6250.

Armenia, J., Wankowicz, S.A.M., Liu, D., Gao, J., Kundra, R., Reznik, E., Chatila, W.K., Chakravarty, D., Han, G.C., Coleman, I., et al. (2018). The long tail of oncogenic drivers in prostate cancer. Nat. Genet. 50, 645–651.

Arora, V.K., Schenkein, E., Murali, R., Subudhi, S.K., Wongvipat, J., Balbas, M.D., Shah, N., Cai, L., Efstathiou, E., Logothetis, C., et al. (2013). Glucocorticoid receptor confers resistance to antiandrogens by bypassing androgen receptor blockade. Cell 155, 1309– 1322.

Augello, M.A., Liu, D., Deonarine, L.D., Robinson, B.D., Huang, D., Stelloo, S., Blattner, M., Doane, A.S., Wong, E.W.P., Chen, Y., et al. (2019). CHD1 Loss Alters AR Binding at Lineage-Specific Enhancers and Modulates Distinct Transcriptional Programs to Drive Prostate Tumorigenesis. Cancer Cell 35, 817–819.

Avsec, Ž., Weilert, M., Shrikumar, A., Krueger, S., Alexandari, A., Dalal, K., Fropf, R., McAnany, C., Gagneur, J., Kundaje, A., et al. Base-resolution models of transcription factor binding reveal soft motif syntax.

Bedford, M.T., and van Helden, P.D. (1987). Hypomethylation of DNA in pathological conditions of the human prostate. Cancer Res. 47, 5274–5276.

Börno, S.T., Fischer, A., Kerick, M., Fälth, M., Laible, M., Brase, J.C., Kuner, R., Dahl, A., Grimm, C., Sayanjali, B., et al. (2012). Genome-wide DNA methylation events in TMPRSS2-ERG fusion-negative prostate cancers implicate an EZH2-dependent mechanism with miR-26a hypermethylation. Cancer Discov. 2, 1024–1035.

Braadland, P.R., and Urbanucci, A. (2019). Chromatin reprogramming as an adaptation mechanism in advanced prostate cancer. Endocr. Relat. Cancer 26, R211–R235.

Buenrostro, J.D., Giresi, P.G., Zaba, L.C., Chang, H.Y., and Greenleaf, W.J. (2013). Transposition of native chromatin for fast and sensitive epigenomic profiling of open chromatin, DNA-binding proteins and nucleosome position. Nat. Methods 10, 1213–1218.

Cancer Genome Atlas Research Network (2015). The Molecular Taxonomy of Primary Prostate Cancer. Cell *163*, 1011–1025.

Chen, Z., Wu, D., Thomas-Ahner, J.M., Lu, C., Zhao, P., Zhang, Q., Geraghty, C., Yan, P.S., Hankey, W., Sunkel, B., et al. (2018). Diverse AR-V7 cistromes in castration-resistant prostate cancer are governed by HoxB13. Proc. Natl. Acad. Sci. U. S. A. 115, 6810–6815.

Corces, M.R., Ryan Corces, M., Granja, J.M., Shams, S., Louie, B.H., Seoane, J.A., Zhou, W., Silva, T.C., Groeneveld, C., Wong, C.K., et al. (2018). The chromatin accessibility landscape of primary human cancers. Science 362, eaav1898.

Eisenberg, E., and Levanon, E.Y. (2013). Human housekeeping genes, revisited. Trends Genet. 29, 569–574.

ENCODE Project Consortium (2012). An integrated encyclopedia of DNA elements in the human genome. Nature 489, 57–74.

Espiritu, S.M.G., Liu, L.Y., Rubanova, Y., Bhandari, V., Holgersen, E.M., Szyca, L.M., Fox, N.S., Chua, M.L.K., Yamaguchi, T.N., Heisler, L.E., et al. (2018). The Evolutionary Landscape of Localized Prostate Cancers Drives Clinical Aggression. Cell 173, 1003– 1013.e15.

Ewing, C.M., Ray, A.M., Lange, E.M., Zuhlke, K.A., Robbins, C.M., Tembe, W.D., Wiley, K.E., Isaacs, S.D., Johng, D., Wang, Y., et al. (2012). Germline mutations in HOXB13 and prostate-cancer risk. N. Engl. J. Med. 366, 141–149.

Faust, G.G., and Hall, I.M. (2014). SAMBLASTER: fast duplicate marking and structural variant read extraction. Bioinformatics 30, 2503–2505.

Fishilevich, S., Nudel, R., Rappaport, N., Hadar, R., Plaschkes, I., Iny Stein, T., Rosen, N., Kohn, A., Twik, M., Safran, M., et al. (2017). GeneHancer: genome-wide integration of enhancers and target genes in GeneCards. Database 2017.

Flavahan, W.A., Gaskell, E., and Bernstein, B.E. (2017). Epigenetic plasticity and the hallmarks of cancer. Science 357.

Friedlander, T.W., Roy, R., Tomlins, S.A., Ngo, V.T., Kobayashi, Y., Azameera, A., Rubin, M.A., Pienta, K.J., Chinnaiyan, A., Ittmann, M.M., et al. (2012). Common structural and epigenetic changes in the genome of castration-resistant prostate cancer. Cancer Res. 72, 616–625.

Gerhauser, C., Favero, F., Risch, T., Simon, R., Feuerbach, L., Assenov, Y., Heckmann, D., Sidiropoulos, N., Waszak, S.M., Hübschmann, D., et al. (2018). Molecular Evolution of Early-Onset Prostate Cancer Identifies Molecular Risk Markers and Clinical Trajectories. Cancer Cell 34, 996–1011.e8.

Gheorghe, M., Sandve, G.K., Khan, A., Chèneby, J., Ballester, B., and Mathelier, A. (2019). A map of direct TF-DNA interactions in the human genome. Nucleic Acids Res. 47, 7715.

Grant, C.E., Bailey, T.L., and Noble, W.S. (2011). FIMO: scanning for occurrences of a given motif. Bioinformatics 27, 1017–1018.

Grasso, C.S., Wu, Y.-M., Robinson, D.R., Cao, X., Dhanasekaran, S.M., Khan, A.P., Quist, M.J., Jing, X., Lonigro, R.J., Brenner, J.C., et al. (2012). The mutational landscape of lethal castration-resistant prostate cancer. Nature 487, 239–243.

Grindstad, T., Andersen, S., Al-Saad, S., Donnem, T., Kiselev, Y., Nordahl Melbø-Jørgensen, C., Skjefstad, K., Busund, L.-T., Bremnes, R.M., and Richardsen, E. (2015). High progesterone receptor expression in prostate cancer is associated with clinical failure. PLoS One 10, e0116691.

Grindstad, T., Richardsen, E., Andersen, S., Skjefstad, K., Rakaee Khanehkenari, M., Donnem, T., Ness, N., Nordby, Y., Bremnes, R.M., Al-Saad, S., et al. (2018). Progesterone Receptors in Prostate Cancer: Progesterone receptor B is the isoform associated with disease progression. Sci. Rep. 8, 11358.

Gundem, G., Van Loo, P., Kremeyer, B., Alexandrov, L.B., Tubio, J.M.C., Papaemmanuil, E., Brewer, D.S., Kallio, H.M.L., Högnäs, G., Annala, M., et al. (2015). The evolutionary history of lethal metastatic prostate cancer. Nature 520, 353–357.

Gupta, S., Stamatoyannopoulos, J.A., Bailey, T.L., and Noble, W. (2007). Quantifying similarity between motifs. Genome Biology 8, R24.

Hankey, W., Chen, Z., and Wang, Q. (2020). Shaping Chromatin States in Prostate Cancer by Pioneer Transcription Factors. Cancer Res. 80, 2427–2436.

Heinz, S., Benner, C., Spann, N., Bertolino, E., Lin, Y.C., Laslo, P., Cheng, J.X., Murre, C., Singh, H., and Glass, C.K. (2010). Simple combinations of lineage-determining transcription factors prime cis-regulatory elements required for macrophage and B cell identities. Mol. Cell 38, 576–589.

Hieronymus, H., Lamb, J., Ross, K.N., Peng, X.P., Clement, C., Rodina, A., Nieto, M., Du, J., Stegmaier, K., Raj, S.M., et al. (2006). Gene expression signature-based chemical genomic prediction identifies a novel class of HSP90 pathway modulators. Cancer Cell 10, 321–330.

Hinrichs, A.S., Karolchik, D., Baertsch, R., Barber, G.P., Bejerano, G., Clawson, H., Diekhans, M., Furey, T.S., Harte, R.A., Hsu, F., et al. (2006). The UCSC Genome Browser Database: update 2006. Nucleic Acids Res. 34, D590–D598.

Isikbay, M., Otto, K., Kregel, S., Kach, J., Cai, Y., Vander Griend, D.J., Conzen, S.D., and Szmulewitz, R.Z. (2014). Glucocorticoid receptor activity contributes to resistance to androgen-targeted therapy in prostate cancer. Horm. Cancer 5, 72–89.

Jiang, L.-H., Zhang, H., and Tang, J.-H. (2018). MiR-30a: A Novel Biomarker and Potential Therapeutic Target for Cancer. J. Oncol. 2018, 5167829.

Jimenez, R.E., Fischer, A.H., Petros, J.A., and Amin, M.B. (2000). Glutathione S-transferase pi gene methylation: the search for a molecular marker of prostatic adenocarcinoma. Adv. Anat. Pathol. 7, 382–389.

Jozwik, K.M., and Carroll, J.S. (2012). Pioneer factors in hormone-dependent cancers. Nat. Rev. Cancer 12, 381–385.

Koh, C.M., Bieberich, C.J., Dang, C.V., Nelson, W.G., Yegnasubramanian, S., and De Marzo, A.M. (2010). MYC and Prostate Cancer. Genes Cancer 1, 617–628.

Kron, K.J., Murison, A., Zhou, S., Huang, V., Yamaguchi, T.N., Shiah, Y.-J., Fraser, M., van der Kwast, T., Boutros, P.C., Bristow, R.G., et al. (2017). TMPRSS2–ERG fusion co-opts master transcription factors and activates NOTCH signaling in primary prostate cancer. Nat. Genet. 49, 1336.

Kulakovskiy, I.V., Vorontsov, I.E., Yevshin, I.S., Sharipov, R.N., Fedorova, A.D., Rumynskiy, E.I., Medvedeva, Y.A., Magana-Mora, A., Bajic, V.B., Papatsenko, D.A., et al. (2018).

HOCOMOCO: towards a complete collection of transcription factor binding models for human and mouse via large-scale ChIP-Seq analysis. Nucleic Acids Res. 46, D252–D259.

Langmead, B., and Salzberg, S.L. (2012). Fast gapped-read alignment with Bowtie 2. Nat. Methods 9, 357–359.

Latonen, L., Afyounian, E., Jylhä, A., Nättinen, J., Aapola, U., Annala, M., Kivinummi, K.K., Tammela, T.T.L., Beuerman, R.W., Uusitalo, H., et al. (2018). Integrative proteomics in prostate cancer uncovers robustness against genomic and transcriptomic aberrations during disease progression. Nat. Commun. 9, 1176.

Lee, W.H., Isaacs, W.B., Bova, G.S., and Nelson, W.G. (1997). CG island methylation changes near the GSTP1 gene in prostatic carcinoma cells detected using the polymerase chain reaction: a new prostate cancer biomarker. Cancer Epidemiol. Biomarkers Prev. 6, 443–450.

Li, H., and Durbin, R. (2009). Fast and accurate short read alignment with Burrows–Wheeler transform. Bioinformatics 25, 1754–1760.

Li, H., Handsaker, B., Wysoker, A., Fennell, T., Ruan, J., Homer, N., Marth, G., Abecasis, G., Durbin, R., and 1000 Genome Project Data Processing Subgroup (2009). The Sequence Alignment/Map format and SAMtools. Bioinformatics 25, 2078–2079.

Losada, A. (2014). Cohesin in cancer: chromosome segregation and beyond. Nat. Rev. Cancer 14, 389–393.

Lupien, M., Eeckhoute, J., Meyer, C.A., Wang, Q., Zhang, Y., Li, W., Carroll, J.S., Liu, X.S., and Brown, M. (2008). FoxA1 translates epigenetic signatures into enhancer-driven lineage-specific transcription. Cell 132, 958–970.

Mahapatra, S., Klee, E.W., Young, C.Y.F., Sun, Z., Jimenez, R.E., Klee, G.G., Tindall, D.J., and Donkena, K.V. (2012). Global methylation profiling for risk prediction of prostate cancer. Clin. Cancer Res. 18, 2882–2895.

Massie, C.E., Lynch, A., Ramos-Montoya, A., Boren, J., Stark, R., Fazli, L., Warren, A., Scott, H., Madhu, B., Sharma, N., et al. (2011). The androgen receptor fuels prostate cancer by regulating central metabolism and biosynthesis. EMBO J. 30, 2719–2733.

Odero-Marah, V., Hawsawi, O., Henderson, V., and Sweeney, J. (2018). Epithelial-Mesenchymal Transition (EMT) and Prostate Cancer. In Cell & Molecular Biology of Prostate Cancer: Updates, Insights and New Frontiers, H. Schatten, ed. (Cham: Springer International Publishing), pp. 101–110.

Parolia, A., Cieslik, M., Chu, S.-C., Xiao, L., Ouchi, T., Zhang, Y., Wang, X., Vats, P., Cao, X., Pitchiaya, S., et al. (2019). Distinct structural classes of activating FOXA1 alterations in advanced prostate cancer. Nature 571, 413–418.

Peng, L., Bian, X.W., Li, D.K., Xu, C., Wang, G.M., Xia, Q.Y., and Xiong, Q. (2015). Large-scale RNA-Seq Transcriptome Analysis of 4043 Cancers and 548 Normal Tissue Controls across 12 TCGA Cancer Types. Sci. Rep. 5, 13413.

Pombo, A., and Dillon, N. (2015). Three-dimensional genome architecture: players and mechanisms. Nat. Rev. Mol. Cell Biol. 16, 245–257.

Pomerantz, M.M., Li, F., Takeda, D.Y., Lenci, R., Chonkar, A., Chabot, M., Cejas, P., Vazquez, F., Cook, J., Shivdasani, R.A., et al. (2015). The androgen receptor cistrome is extensively reprogrammed in human prostate tumorigenesis. Nat. Genet. 47, 1346–1351.

Pomerantz, M.M., Qiu, X., Zhu, Y., Takeda, D.Y., Pan, W., Baca, S.C., Gusev, A., Korthauer, K.D., Severson, T.M., Ha, G., et al. (2020). Prostate cancer reactivates developmental epigenomic programs during metastatic progression. Nat. Genet.

Quigley, D.A., Dang, H.X., Zhao, S.G., Lloyd, P., Aggarwal, R., Alumkal, J.J., Foye, A., Kothari, V., Perry, M.D., Bailey, A.M., et al. (2018). Genomic Hallmarks and Structural Variation in Metastatic Prostate Cancer. Cell 175, 889.

Quinlan, A.R., and Hall, I.M. (2010). BEDTools: a flexible suite of utilities for comparing genomic features. Bioinformatics 26, 841–842.

Rajbhandari, P., Thomas, B.J., Feng, A.-C., Hong, C., Wang, J., Vergnes, L., Sallam, T., Wang, B., Sandhu, J., Seldin, M.M., et al. (2018). IL-10 Signaling Remodels Adipose Chromatin Architecture to Limit Thermogenesis and Energy Expenditure. Cell 172, 218– 233.e17.

Rickman, D.S., Soong, T.D., Moss, B., Mosquera, J.M., Dlabal, J., Terry, S., MacDonald, T.Y., Tripodi, J., Bunting, K., Najfeld, V., et al. (2012). Oncogene-mediated alterations in chromatin conformation. Proc. Natl. Acad. Sci. U. S. A. 109, 9083–9088.

Roadmap Epigenomics Consortium, Kundaje, A., Meuleman, W., Ernst, J., Bilenky, M., Yen, A., Heravi-Moussavi, A., Kheradpour, P., Zhang, Z., Wang, J., et al. (2015). Integrative analysis of 111 reference human epigenomes. Nature 518, 317–330.

Robinson, D., Van Allen, E.M., Wu, Y.-M., Schultz, N., Lonigro, R.J., Mosquera, J.-M., Montgomery, B., Taplin, M.-E., Pritchard, C.C., Attard, G., et al. (2015). Integrative clinical genomics of advanced prostate cancer. Cell 161, 1215–1228.

Rodriguez-Bravo, V., Carceles-Cordon, M., Hoshida, Y., Cordon-Cardo, C., Galsky, M.D., and Domingo-Domenech, J. (2017). The role of GATA2 in lethal prostate cancer aggressiveness. Nat. Rev. Urol. 14, 38–48.

Rowley, M.J., Jordan Rowley, M., and Corces, V.G. (2018). Organizational principles of 3D genome architecture. Nature Reviews Genetics 19, 789–800.

Sahu, B., Laakso, M., Ovaska, K., Mirtti, T., Lundin, J., Rannikko, A., Sankila, A., Turunen, J.-P., Lundin, M., Konsti, J., et al. (2011). Dual role of FoxA1 in androgen receptor binding to chromatin, androgen signalling and prostate cancer. EMBO J. 30, 3962–3976.

Sandoval, G.J., Pulice, J.L., Pakula, H., Schenone, M., Takeda, D.Y., Pop, M., Boulay, G., Williamson, K.E., McBride, M.J., Pan, J., et al. (2018). Binding of TMPRSS2-ERG to BAF Chromatin Remodeling Complexes Mediates Prostate Oncogenesis. Mol. Cell 71, 554– 566.e7.

Scharer, C.D., Barwick, B.G., Guo, M., Bally, A.P.R., and Boss, J.M. (2018). Plasma cell differentiation is controlled by multiple cell division-coupled epigenetic programs. Nat. Commun. 9, 1698.

Sharma, N.L., Massie, C.E., Ramos-Montoya, A., Zecchini, V., Scott, H.E., Lamb, A.D., MacArthur, S., Stark, R., Warren, A.Y., Mills, I.G., et al. (2013). The androgen receptor induces a distinct transcriptional program in castration-resistant prostate cancer in man. Cancer Cell 23, 35–47.

Siegel, R.L., Miller, K.D., and Jemal, A. (2018). Cancer statistics, 2018. CA: A Cancer Journal for Clinicians 68, 7–30.

Sinha, A., Huang, V., Livingstone, J., Wang, J., Fox, N.S., Kurganovs, N., Ignatchenko, V., Fritsch, K., Donmez, N., Heisler, L.E., et al. (2019). The Proteogenomic Landscape of Curable Prostate Cancer. Cancer Cell 35, 414–427.e6.

Stelloo, S., Nevedomskaya, E., Kim, Y., Schuurman, K., Valle-Encinas, E., Lobo, J., Krijgsman, O., Peeper, D.S., Chang, S.L., Feng, F.Y.-C., et al. (2018). Integrative epigenetic taxonomy of primary prostate cancer. Nat. Commun. 9, 4900.

Taberlay, P.C., Achinger-Kawecka, J., Lun, A.T.L., Buske, F.A., Sabir, K., Gould, C.M., Zotenko, E., Bert, S.A., Giles, K.A., Bauer, D.C., et al. (2016). Three-dimensional disorganization of the cancer genome occurs coincident with long-range genetic and epigenetic alterations. Genome Res. 26, 719–731.

Takeda, D.Y., Spisák, S., Seo, J.-H., Bell, C., O’Connor, E., Korthauer, K., Ribli, D., Csabai, I., Solymosi, N., Szállási, Z., et al. (2018). A Somatically Acquired Enhancer of the Androgen Receptor Is a Noncoding Driver in Advanced Prostate Cancer. Cell 174, 422–432.e13.

Thurman, R.E., Rynes, E., Humbert, R., Vierstra, J., Maurano, M.T., Haugen, E., Sheffield, N.C., Stergachis, A.B., Wang, H., Vernot, B., et al. (2012). The accessible chromatin landscape of the human genome. Nature 489, 75–82.

Toenhake, C.G., Fraschka, S.A.-K., Vijayabaskar, M.S., Westhead, D.R., van Heeringen, S.J., and Bártfai, R. (2018). Chromatin Accessibility-Based Characterization of the Gene Regulatory Network Underlying Plasmodium falciparum Blood-Stage Development. Cell Host Microbe 23, 557–569.e9.

Urbanucci, A., Sahu, B., Seppälä, J., Larjo, A., Latonen, L.M., Waltering, K.K., Tammela, T.L.J., Vessella, R.L., Lähdesmäki, H., Jänne, O.A., et al. (2012). Overexpression of androgen receptor enhances the binding of the receptor to the chromatin in prostate cancer. Oncogene 31, 2153–2163.

Urbanucci, A., Barfeld, S.J., Kytölä, V., Itkonen, H.M., Coleman, I.M., Vodák, D., Sjöblom, L., Sheng, X., Tolonen, T., Minner, S., et al. (2017). Androgen Receptor Deregulation Drives Bromodomain-Mediated Chromatin Alterations in Prostate Cancer. Cell Rep. 19, 2045–2059.

Varambally, S., Dhanasekaran, S.M., Zhou, M., Barrette, T.R., Kumar-Sinha, C., Sanda, M.G., Ghosh, D., Pienta, K.J., Sewalt, R.G.A.B., Otte, A.P., et al. (2002). The polycomb group protein EZH2 is involved in progression of prostate cancer. Nature 419, 624–629.

Viré, E., Brenner, C., Deplus, R., Blanchon, L., Fraga, M., Didelot, C., Morey, L., Van Eynde, A., Bernard, D., Vanderwinden, J.-M., et al. (2006). The Polycomb group protein EZH2 directly controls DNA methylation. Nature 439, 871–874.

Viswanathan, S.R., Ha, G., Hoff, A.M., Wala, J.A., Carrot-Zhang, J., Whelan, C.W., Haradhvala, N.J., Freeman, S.S., Reed, S.C., Rhoades, J., et al. (2018). Structural Alterations Driving Castration-Resistant Prostate Cancer Revealed by Linked-Read Genome Sequencing. Cell 174, 433–447.e19.

Wang, Q., Li, W., Zhang, Y., Yuan, X., Xu, K., Yu, J., Chen, Z., Beroukhim, R., Wang, H., Lupien, M., et al. (2009). Androgen receptor regulates a distinct transcription program in androgen-independent prostate cancer. Cell 138, 245–256.

Wang, Y., Song, F., Zhang, B., Zhang, L., Xu, J., Kuang, D., Li, D., Choudhary, M.N.K., Li, Y., Hu, M., et al. (2018). The 3D Genome Browser: a web-based browser for visualizing 3D genome organization and long-range chromatin interactions. Genome Biology 19.

Weischenfeldt, J., Dubash, T., Drainas, A.P., Mardin, B.R., Chen, Y., Stütz, A.M., Waszak, S.M., Bosco, G., Halvorsen, A.R., Raeder, B., et al. (2017). Pan-cancer analysis of somatic copy-number alterations implicates IRS4 and IGF2 in enhancer hijacking. Nat. Genet. 49, 65–74.

Wu, J., Xu, J., Liu, B., Yao, G., Wang, P., Lin, Z., Huang, B., Wang, X., Li, T., Shi, S., et al. (2018). Chromatin analysis in human early development reveals epigenetic transition during ZGA. Nature 557, 256–260.

Xu, K., Wu, Z.J., Groner, A.C., He, H.H., Cai, C., Lis, R.T., Wu, X., Stack, E.C., Loda, M., Liu, T., et al. (2012). EZH2 oncogenic activity in castration-resistant prostate cancer cells is Polycomb-independent. Science 338, 1465–1469.

Yevshin, I., Sharipov, R., Kolmykov, S., Kondrakhin, Y., and Kolpakov, F. (2019). GTRD: a database on gene transcription regulation—2019 update. Nucleic Acids Res. 47, D100– D105.

Ylipää, A., Kivinummi, K., Kohvakka, A., Annala, M., Latonen, L., Scaravilli, M., Kartasalo, K., Leppänen, S.-P., Karakurt, S., Seppälä, J., et al. (2015). Transcriptome Sequencing Reveals PCAT5 as a Novel ERG-Regulated Long Noncoding RNA in Prostate Cancer. Cancer Res. 75, 4026–4031.

Yu, J., Yu, J., Mani, R.-S., Cao, Q., Brenner, C.J., Cao, X., Wang, X., Wu, L., Li, J., Hu, M., et al. (2010). An integrated network of androgen receptor, polycomb, and TMPRSS2-ERG gene fusions in prostate cancer progression. Cancer Cell 17, 443–454.

Zhang, Y., Liu, T., Meyer, C.A., Eeckhoute, J., Johnson, D.S., Bernstein, B.E., Nusbaum, C., Myers, R.M., Brown, M., Li, W., et al. (2008). Model-based analysis of ChIP-Seq (MACS). Genome Biol. 9, R137.

Zhao, S.G., Chen, W.S., Li, H., Foye, A., Zhang, M., Sjöström, M., Aggarwal, R., Playdle, D., Liao, A., Alumkal, J.J., et al. (2020). The DNA methylation landscape of advanced prostate cancer. Nat. Genet. 52, 778–789.

Zhou, Y., Huang, T., Cheng, A.S.L., Yu, J., Kang, W., and To, K.F. (2016). The TEAD Family and Its Oncogenic Role in Promoting Tumorigenesis. Int. J. Mol. Sci. 17, 138.

